# Overcoming constraints on the detection of recessive selection in human genes from population frequency data

**DOI:** 10.1101/2021.05.06.443024

**Authors:** Daniel J. Balick, Daniel M. Jordan, Shamil Sunyaev, Ron Do

**Affiliations:** Division of Genetics, Brigham and Women’s Hospital, Harvard Medical School, Boston, MA, USA; Department of Biomedical Informatics, Harvard Medical School, Boston, MA, USA; The Charles Bronfman Institute for Personalized Medicine, Icahn School of Medicine at Mount Sinai, New York, NY, USA; Department of Genetics and Genomic Sciences, Icahn School of Medicine at Mount Sinai, New York, NY, USA

## Abstract

The identification of genes that evolve under recessive natural selection is a longstanding goal of population genetics research with important applications to disease gene discovery. We found that commonly used methods to evaluate selective constraint at the gene level are highly sensitive to genes under heterozygous selection but ubiquitously fail to detect recessively evolving genes. Additionally, more sophisticated likelihood-based methods designed to detect recessivity similarly lack power for a human gene of realistic length from current population sample sizes. However, extensive simulations suggested that recessive genes may be detectable in aggregate. Here, we offer a method informed by population genetics simulations designed to detect recessive purifying selection in gene sets. Applying this to empirical gene sets produced significant enrichments for strong recessive selection in genes previously inferred to be under recessive selection in a consanguineous cohort and in genes involved in autosomal recessive monogenic disorders.

## Introduction

Identifying human genes that evolve under recessive natural selection remains incredibly difficult, despite a host of efforts to characterize the dominance of human traits and diseases [1–7]. Substantial progress has been made to quantify features of the joint distribution of dominance and selection coefficients in model organisms like drosophila and yeast [8–10]. However, in humans, as with most long-lived organisms, direct experimentation is not possible, so the search for recessive selection relies largely on inference from natural population data. Previous efforts span from the sequencing of consanguineous cohorts to inference by analogy to experimental systems [11–15]. However, with the advent of large-scale sequence data, it may be possible to identify at least some subset of recessive natural selection using information from observables like features of the site frequency spectrum [16–18].

Many analytic and computational tools have been developed to identify natural selection, but in general their effectiveness is dependent on selection in heterozygous form [19–22]. This is primarily due to the fact that purifying selection drives variants to low frequencies, making the probability of randomly forming homozygotes (or compound heterozygotes with similar function) vanishingly small. Recessive selection is complicated by the fact that it appears effectively neutral at these low frequencies since the formation of homozygotes is improbable, precluding efficient purifying selection on homozygotes. Thus, the essential problem in identifying recessive deleterious variation is not disentangling it from additive or dominant variation, but rather from neutrality and the genomic background at large. This also explains why most tools designed to probe purifying selection, while highly successful at identifying variation when additive or dominant, are largely insensitive to recessivity, conflating it with neutral variation or weakly deleterious additivity [23]. Notably, despite proposed methods to infer these parameters from population frequency data [17], diploid selection coefficients are not yet known for individual genes in the human genome.

## Results

### Insensitivity of existing statistics to recessive variation

In a recent perspective by Fuller, et al., the authors demonstrated that the probability of Loss-of-function Intolerance (pLI) score [21], one of the most commonly used scores of selection in humans, is well powered to identify heterozygous selection but is largely insensitive to recessive variation [23]. This score identifies constrained regions by detecting the reduction of frequency or absence of variation in a large human population sample. However, recessive selection is often undetectable by this measure, as variation at extremely low frequencies is necessarily in heterozygote form preventing the action of selection on homozygotes. This highlights the central problem in identifying recessive variation: diploidy can fundamentally shield mutations from homozygous selection.

To expand on this observation, we compiled a literature-based gene set expected to be enriched for recessive selection (n=606) from genes identified in a cohort of British individuals of Pakistani descent exhibiting a high degree of consanguinity (henceforth “ConsangBP”) [11]. These genes were identified as genes harboring rare homozygous Loss of Function (LOF) variants and were measured in aggregate to be under significant recessive purifying selection. We assembled an analogous gene set expected to be depleted for recessive selection (n=825) from an empirically derived probability of haploinsufficiency (*p*_*HI*_), restricting to genes with *p*_*HI*_ ≥ 0.8 (henceforth “HI80”) [24]. Using these two gene sets, we tested the ability of several commonly used measures of selection to differentiate either recessive or non-recessive genes from the rest of the genome. For consistency with later analyses, we restricted both sets to genes with aggregated LOF and PolyPhen2 probably damaging (henceforth “damaging”) mutation rates above 3 × 10^−6^ per haploid individual per generation (ConsangBP n =384 genes, HI80 n=574, whole genome n=8316) [25].

We tested a variety of commonly applied per gene statistics that rely on a combination of intra-species population genetic information, inter-species comparative genomics, and functional information to score each gene (see **Figure 1**) [19–22, 26–29]. All population genetic values were computed using data from the non-Finnish European (NFE) cohort of the Exome Aggregation Consortium (ExAC) for consistency with later analyses [21]. Nearly all statistics shown are highly sensitive to purifying selection in genes under non-recessive selection, but all were generally insensitive to genes under recessive selection. The net impact of this insensitivity on genome-wide properties of natural selection is unclear, as the number of genes that behave recessively remains unmeasured in humans. Since such measures are frequently used to prioritize genes for clinical and biological studies, this has great potential impact across both biology and medicine.

**Figure 1:**
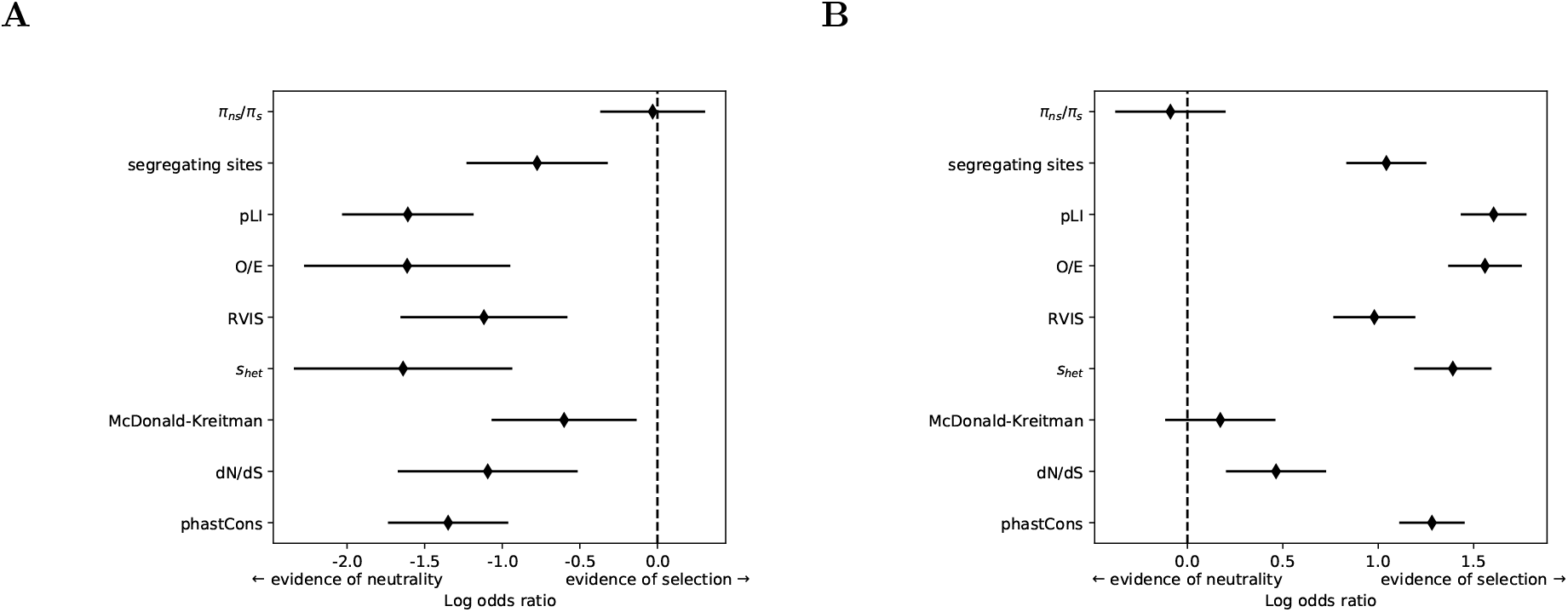
*Enrichment for genes under selection using various per gene metrics*. Points show enrichment for genes showing evidence of selection according to each per gene metric, expressed as log odds ratio. Lines show 95% confidence intervals. The metrics shown include scores based on population constraint within humans (ratio of nonsynonymous to synonymous nucleotide diversity *π*_*ns*_*/π*_*s*_, number of nonsynonymous segregating sites, and constraint scores PLI [21], OE [28], RVIS [27], and S_*het*_ [29]), scores based on conservation between species (dN/dS score [19, 20], phastCons conservation score [22]), and a hybrid score that includes both (McDonald-Kreitman neutrality index [26]). For details of how these scores were processed, see Methods. **A**. The putatively recessive ConsangBP gene set [11] showed either a depletion for genes with evidence of selection or a lack of power in all scores tested. **B**. The putatively non-recessive HI80 gene set [24] showed either an enrichment for genes with evidence of selection or a lack of power in all scores tested.

### Power to infer recessive selection at the single gene level

It is unclear whether the insensitivity of common measures to recessive selection is due to features of these specific statistics or a fundamental power limitation. A partial answer was provided by Williamson, et al. who developed a Poisson Random Field (PRF) based log likelihood ratio test (LLRT) to evaluate dominance using population genetic data [17, 30]. This test compares the null hypothesis of additive selection (*h* = 0.5) to an alternative hypothesis of unrestricted dominance (0 ≤ *h* ≤ 0.5). They demonstrate that this test has ample power to reject the null hypothesis when selection is recessive if there are a sufficient number of segregating sites. However, they focused on the ability to reject the null hypothesis of uniformly additive selection across the entire genome using data available in 2004, and this analysis does not address whether modern population genomics datasets can be used to detect recessive selection on the level of a single gene. Additionally, they assumed an equilibrium demography rather than a more realistic demography for human populations, which becomes more important with increasing sample size.

We simulated alleles for a grid of selection and dominance coefficients through a realistic demography, and downsampled to the size of the NFE subsample of the ExAC dataset, the largest relatively panmictic single ancestry cohort without recurrent non-CpG mutations [21]. This was repeated for a range of mutational target sizes, representing the combined target size for LOF and damaging missense mutations for human genes. We focused on recessive simulations under strong selection (*h* = 0, *s* = −0.1) to assess the ability of the proposed test to reject the null hypothesis of additivity. As shown in **Figure 2A**, the power to reject the null hypothesis was essentially zero for any realistic gene length (also see Methods and **Supplementary Figures SI1 and SI2**). This is due to ambiguity in the SFS between strong recessive selection and weaker additive selection when there are an insufficient number of segregating sites, which makes it difficult to confidently reject the null hypothesis when selection is truly recessive (*h* = 0) (see **Supplementary Figure SI4**). Our results indicate that this ambiguity is a fundamental feature of recessive selection that exists even under ideal circumstances when demography and mutation rates are known exactly, and is limited in power by finite gene size. These results are entirely consistent with the observations of Fuller et al. [23], as well as with Williamson et al. [17], who estimated the number of segregating sites required to confidently reject the null hypothesis of additivity at a much higher number than the target size for all LOF and damaging mutations in a typical gene (shown in **Supplemental Figure SI1**).

**Figure 2:**
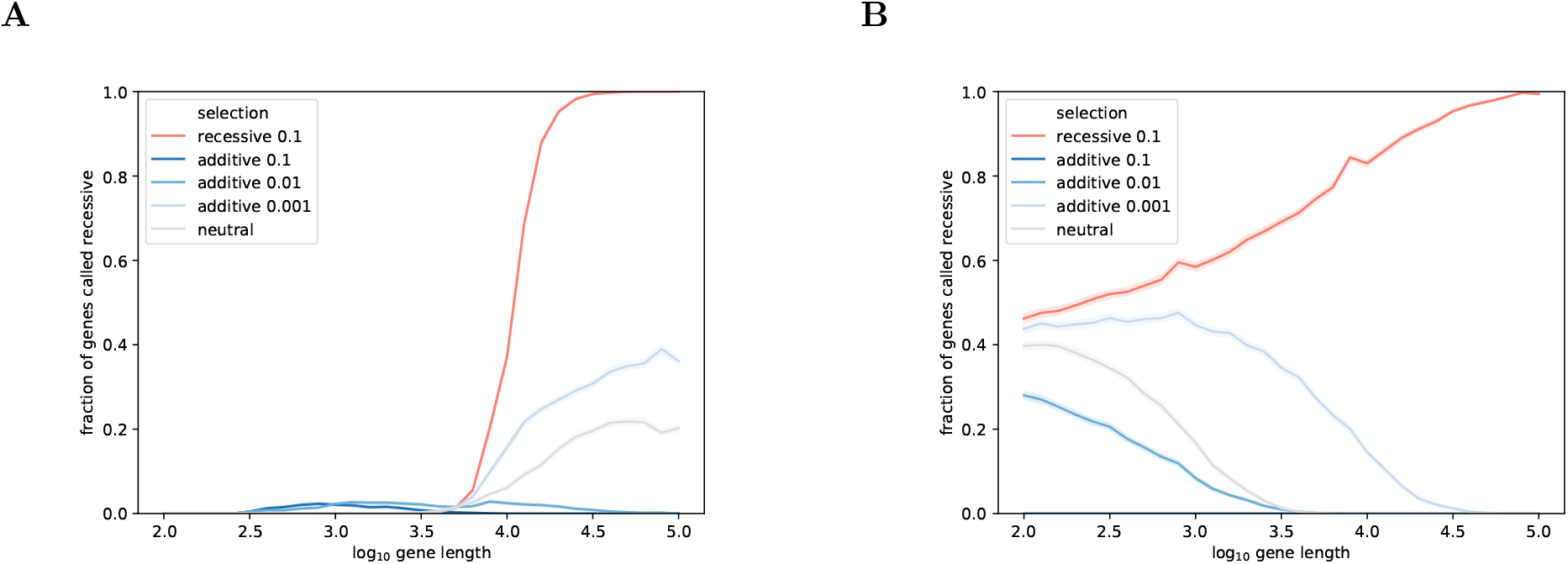
*Power to correctly predict strong recessive selection in a single gene as a function of gene length*. Power to correctly identify simulated genes under strong recessive selection (h=0, s=-0.1; red) is plotted as a function of gene length. False positive rates are shown for genes under additive selection with varying selection strengths: strong selection (s=-0.1, dark blue), weak selection (s=-0.01 and -0.001, lighter blues), and neutrality (s=0, grey). For simplicity, all sites within each gene are assumed to have uniform dominance and selection coefficients, which may be a reasonable approximation within a single functional class (e.g. synonymous sites only, damaging sites only). n=1000 replicate simulations were performed using a demography inferred from the ExAC NFE sample [29, 31]. **A**. Power and false positive rates using the log likelihood ratio test to reject additivity of all selection strengths. Roughly 10,000 sites (corresponding to a mutational target size of 10^−4^) are needed to gain sufficient power. Virtually no genes in the human genome have LOF and damaging target sizes on this order (see **Figure SI1**). **B**. Power and false positive rates using the srML test with the same likelihood function. Some power can be seen starting at roughly 300 sites (mutational target size of 3 *×* 10^−6^). However, false positives persist at high fractions, particularly for additive variation under weak selection. Precise error rates for a set of genes depend both on the length distribution and on the distribution of diploid selection coefficients.

We next considered whether a hypothesis test aimed more narrowly at identifying strong recessive selection might have more power for this task under the same ideal circumstances. We performed an alternative test comparing strong recessive selection (*h <* 0.1, *s* ≥ −0.1) to a model that excludes strong recessive selection (0.5 ≥ *h >* 0 and/or 0 ≥ *s >* −0.1). We identify genes as “strong recessive” if the values of *h* and *s* that maximize likelihood lie within the strong recessive region, and as “non-strong-recessive” otherwise. This test, which we refer to as a strong recessive maximum likelihood (srML) test, also had low power to correctly estimate selection and dominance coefficients for genes under strong recessive selection (see **Figure 2B**). This implies that even with a test specifically designed to maximize power to detect strong recessive selection, detecting individual genes under strong recessive selection based solely on population genetics remains infeasible with current data.

### Power to detect enrichment of recessive selection in gene sets

Given the failure to identify single recessive genes, we next asked if such genes could be identified in aggregate in the form of gene set enrichment over the genomic background. To test this, we simulated gene sets with a range of odds ratios for strong recessive genes to assess the power to detect enrichment over a simulated genomic background.

As described in Methods, we simulated a range of realistic human genomic backgrounds consistent with previous studies and parameterized by the fraction of the genes evolving under strong recessive selection, *f*_*R*_, up to a maximum of 30%. We then simulated gene sets of various sizes (n= {0,100,300,…}) with a known odds ratio from a depletion of OR=0.25 to an enrichment of OR=4.0 for recessive strong selection genes with the remaining genes randomly pulled from a genomic background with a given fraction *f*_*R*_. Both the backgrounds and test sets were sampled from the empirical distribution of ExAC NFE LOF and damaging mutation rates. Using simulated genomic backgrounds, we assessed the power to detect enrichments for the two methods described above. We found no significant enrichment for any plausible gene set size when using the LLRT for rejection of additivity, independent of the odds ratio in the test gene set (**Supplementary Figure SI5**). This is, in part, due to the nature of the test, as recessive genes and weak additive genes have similar PRF likelihood values that make it difficult to confidently reject additivity.

In contrast, we found that gene sets containing a sufficient odds ratio for strong recessive genes can be significantly enriched for genes identified as strong recessive by the srML test. **Figure 3A** shows the power profile for an example simulated genome in which 10% of all genes are under strong recessive selection (*f*_*R*_ = 0.1). The statistical significance is shown as a function of the input odds ratio for various simulated set sizes ranging from 30 to 3000 genes. For relatively small gene sets on the order of 30–100 genes, only odds ratios with more than roughly threefold enrichment show significance, while much larger sets on the order of 1000–3000 genes become significant at more modest values around a 1.3-fold odds ratio. As a point of comparison, **Figure 3B** shows the same plot for a simulated genome in which only 3% of genes are under strong recessive selection (*f*_*R*_ = 0.03). The qualitative features of this power plot are the same when rescaling the horizontal axis, but the significance becomes much weaker, with roughly a 2.5-fold enrichment needed even for gene sets of order 1000. For a full range of simulated genomes with distinct fractions of strong recessivity, see **Supplementary Figure SI6**. See **Supplementary Figure SI7** for the effect of varying *f*_*R*_ for a fixed gene set size. Together, these plots show that the quantitative requirement for a statistically observable enrichment depends sensitively on the fraction of the genome as a whole that is under strong recessive purifying selection—a completely unknown parameter in humans. To expand on this, we plotted the relationship of the estimated odds ratio (using srML on scored genes) on the simulated odds ratio. **Figure 3C** shows how the estimated OR deviates from the true OR as a function of the fraction of the background under strong recessive selection. The odds ratio is an underestimate of the true enrichment (or depletion), making it impossible to infer the number of strong recessive true positives without knowing the exact content of the genomic background at large (i.e. the joint distribution of dominance and selection coefficients). Therefore, an observation of significant enrichment for strong recessive selection likely indicates a true signal but says little about the number of genes driving this signal without additional information.

**Figure 3:**
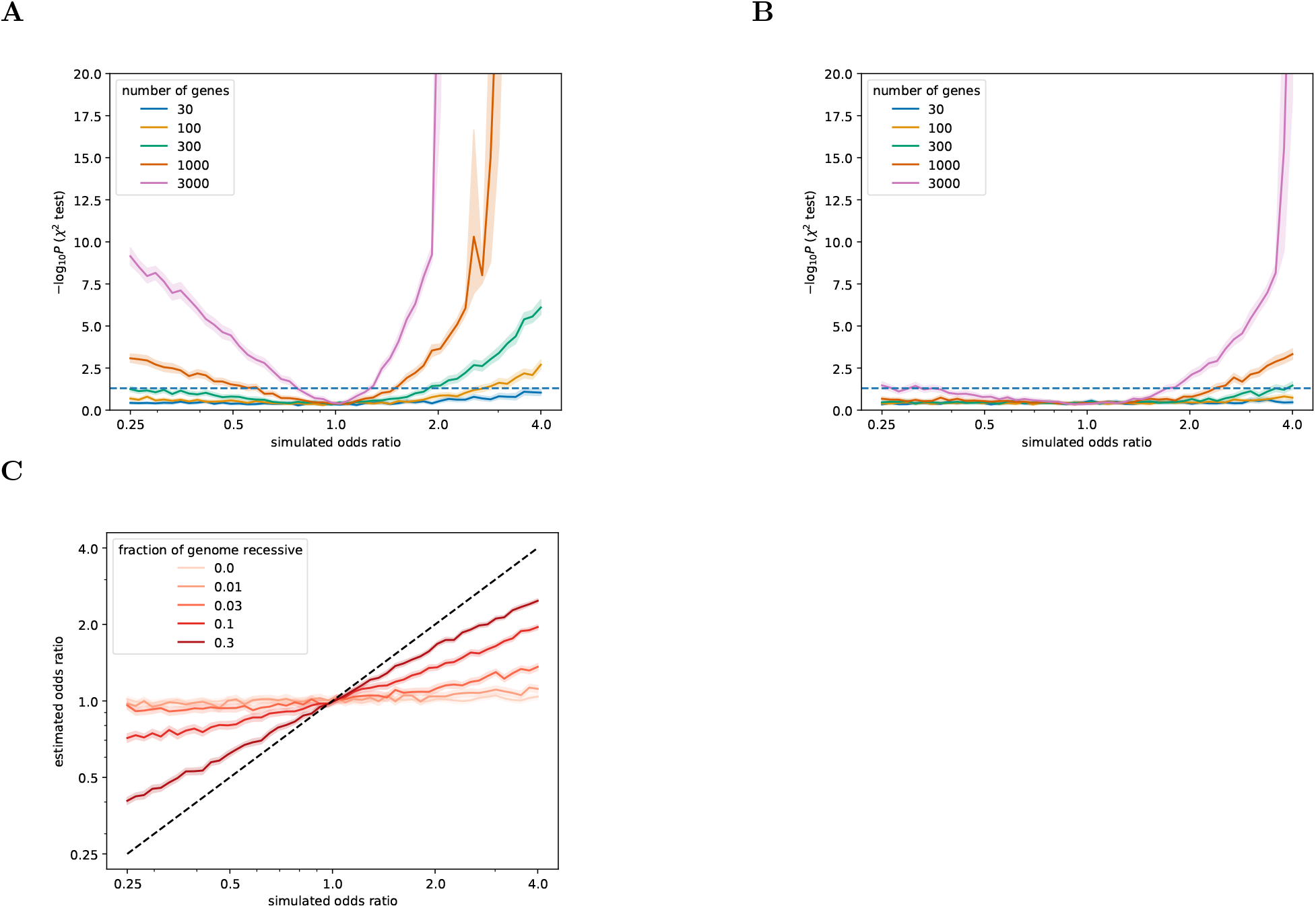
*Power to detect enrichment for genes under strong recessive selection*. Power of the srML test to detect the enrichment of genes under strong recessive selection is plotted as a function of gene set size and true odds ratio of the gene set for two different values of the parameter *f*_*R*_, representing the fraction of the genome that is under strong recessive selection. Qualitative features do not depend on the genome-wide prevalence of strong recessive selection, but the significance threshold is highly sensitive to this parameter. In all cases, the srML test was used to score individual genes, and enrichment of genes predicted recessive was evaluated using a chi-squared contingency test. **A**. Power to detect enrichment of recessive selection for gene sets of various sizes for a simulated genome with 10% of genes under strong recessive selection. **B**. The comparable power plot for a simulated genome with 3% of genes under strong recessive selection. **C**. Estimated odds ratio vs simulated odds ratio for gene sets of size n=300 with varying fractions of the genome under strong recessive selection. srML universally underestimates the true enrichment (slope *<* 1) but is a better estimate for larger recessive fractions of the background (an unknown quantity in humans). Gene set size affects the variance of this dependence, but not the slope.

### Application to empirical gene sets

With a simulation-based understanding of the power to detect enrichment for strong recessive genes in sufficiently large gene sets, we applied the same methodology to gene sets of interest using ExAC NFE data [21]. We assembled several literature-based gene sets with reasonable expectation to be either enriched or depleted for recessive genes under strong selection. Using simulations of a range of dominance and selection coefficients matched to each gene length, we used the PRF-based likelihood to estimate the values of *h* and *s* associated with each gene maximized over 17 possible *h* and *s* pairs (see Methods and **Supplementary File 1** for a list of target sizes and ML *h* and *s* values for all genes). Genes with LOF and damaging mutational target sizes below 3 × 10^−6^ (average of less than 300 sites) were removed from all lists and the genomic background, as shorter genes showed no power to convey meaningful information in simulations (see **Figure 2B**). This provided a total of 8316 scored genes that comprise roughly 40% of the whole genome we were able to annotate. We performed the srML test described above on each gene and tested each gene set against the rest of the genome for enrichment of specific selection classes. Odds ratios and log p-values for enrichment at two primary diploid selection strengths of interest—strong recessive selection and strong additive selection—are summarized in **Figure 4**, along with the number of scored genes in each set.

**Figure 4:**
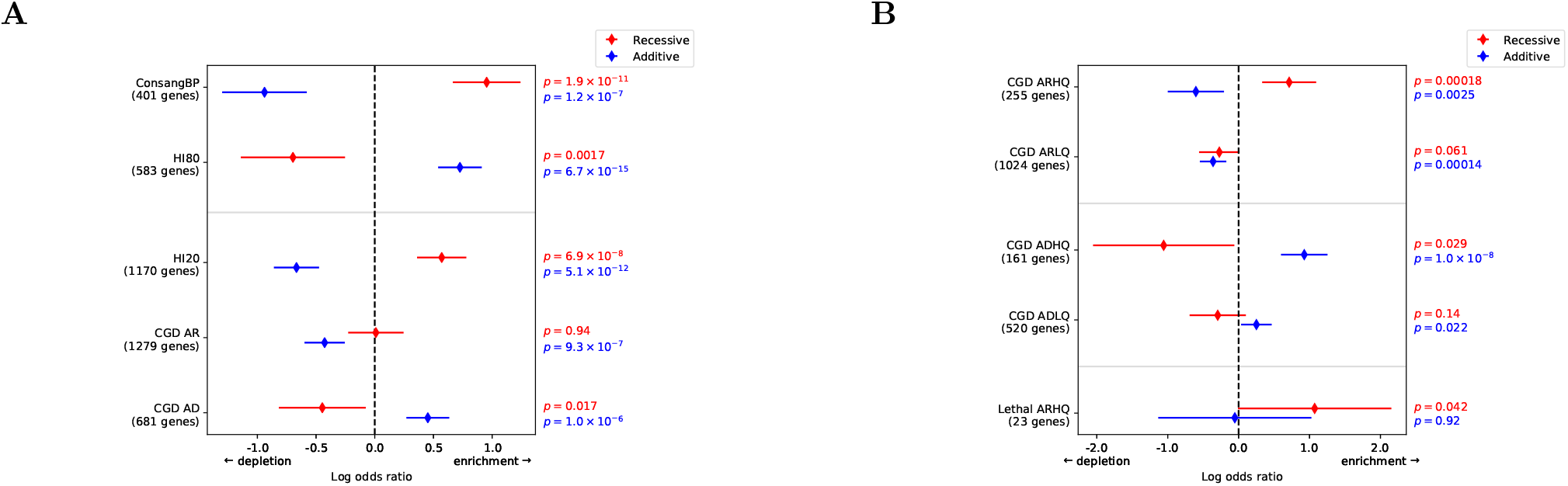
*Enrichment of literature-based gene sets for genes under strong recessive or strong additive natural selection*. Log odds ratio of enrichment for genes called “strong recessive” by the srML test or genes called strong additive by the analogous test for strong additive selection for various gene sets. **A**. ConsangBP contains genes with evidence of recessive selection from a consanguineous British Pakistani population [11]; HI80 contains genes with evidence of haploinsufficiency from an empirically derived haploinsufficiency score [24]; HI20 contains genes with evidence against haploinsufficiency from the same empirically derived haploinsufficiency score; CGD AR and AD contain genes known to be implicated in autosomal recessive or autosomal dominant disease, respectively [7]. **B**. CGD ARHQ and ADHQ sets consist of the subset of CGD AR or AD genes [7] harboring more than one variant annotated with a quality score of 2 stars (“multiple submitters, no conflicts”) or higher in ClinVar [32]; CGD ARLQ and ADLQ consist of all genes in CGD AR or AD that do not meet this criterion; Lethal ARHQ contains genes that are both confidently identified as causal for effectively lethal AR disease and pass the ClinVar quality filter [32, 33].

We analyzed the ConsangBP and HI80 gene sets described above as respective positive and negative controls for the enrichment of genes evolving under strong recessive selection. Enrichment results for both gene sets are shown in **Figure 4A** and detailed in **Supplementary Table S1**. The putatively recessive-enriched ConsangBP set showed a highly significant enrichment for strong recessive selection (OR=2.60, p=1.9 × 10^−11^), as well as a highly significant depletion for strong additive selection (OR=0.50, p=1.2 × 10^−7^). A priori, it is possible to find enrichments in both bins simultaneously for a given gene set, as enrichment for strong recessive genes does not necessitate depletion for strong additive genes, and vice versa. However, our simulations demonstrated that for a homogenous set of strong recessive genes, misclassification as strong additive is unlikely (see **Figure 2B**), suggesting that the ConsangBP gene set contains a large number of strong recessive genes and few strong additive genes compared to the whole genome. The putatively recessive-depleted HI80 gene set showed the opposite behavior, with significant depletion of strong recessive genes (OR=0.5, p=0.002) and highly significant enrichment for strong additive genes (OR=2.07, p=6.7 × 10^−15^). This expected result provides a negative control in which genes identified for their additive or dominant mechanism appear underrepresented for those under strong recessive purifying selection. The depletion of recessive genes in HI80 is less significant than the enrichment of ConsangBP, consistent with the power analysis from simulations (see **Figure 3**). As an additional control, we restricted both gene sets to synonymous variants and found no significant difference from the whole genome, suggesting no major effects due to linkage on the genic level. Last, we confirmed that potential length bias had no significant effect on our enrichments (see Methods and **Supplementary Table S2**).

Having validated the ability to identify gene sets enriched or depleted for strong recessive selection using the ConsangBP and HI80 controls, we applied the same methodology to several additional gene sets. First, we analyzed genes with a low probability (*p* ≤ 0.2) of haploinsufficiency (referred to as HI20) using the same HI predictions used for the HI80 gene set [24]. A priori, it is unclear whether the lack of evidence for haploinsufficient activity is predictive of enrichment over the genomic background for genes under strong recessive selection. We found a qualitatively similar enrichment pattern to that of the recessive control with highly significant enrichment for strong recessive (OR=1.77, p=7.0 × 10^−8^) and depletion of strong additive genes (OR=0.51, p=5.1 × 10^−12^) (see **Figure 4A**). This suggests that the lack of evidence for haploinsufficiency represented by low predicted HI probability is indeed indicative of the presence of genes under strong recessive selection, rather than simply the remainder of genes outside of the strong additive corner. This provides a biological example of the utility of this method to identify correlations to strong recessive selection in aggregate.

Next, we focused on gene sets assembled from data on the mode of inheritance (MOI) of human disease phenotypes. We used annotations from the Clinical Genomic Database (CGD) to create a list of autosomal recessive (AR) disease genes (i.e. genes only containing known AR disease variants), and a list of autosomal dominant (AD) disease genes (those only containing known AD disease variants) [7]. See Methods for details. We assessed the presence of strong recessive genes for each MOI separately by scoring all genes of sufficient length and computing enrichment odds ratios and p-values (see **Figure 4A**). CGD AD genes were depleted for strong recessive selection (OR=0.64, p=0.017) and highly enriched for strong additive selection (OR=1.57, p=1.0 × 10^−6^), qualitatively consistent with the HI80 enrichment profile for non-additive genes but with reduced significance. In contrast, the set of CGD AR genes showed no significant enrichment for genes under strong recessive selection (OR=1.01, p=0.94), despite highly significant depletion for genes under strong additive selection (OR=0.65, p=9.3 × 10^−7^). We hypothesized that this was due to annotation quality for recessive disease genes, some of which might be attributable to the insensitivity of commonly used prioritization statistics and annotation tools to recessive selection (see **Figure 1**). To address this, we applied a ClinVar quality filter to restrict to genes with multiple high quality variant annotations, as detailed in Methods [32]. We stratified the AR list into high quality (CGD ARHQ) and low quality (CGD ARLQ) subsets, with the latter containing all genes failing our quality filtration step. We analogously stratified AD into CGD ADHQ and CGD ADLQ for comparison. The resulting enrichments are displayed in **Figure 4B**. ARHQ showed a significant enrichment for strong recessive genes (OR=2.04, p=1.8 × 10^−4^) and significant depletion for strong additive genes (OR=0.55, p=2.5 × 10^−3^), consistent with naïve expectations for genes responsible for, often severe, autosomal recessive Mendelian disease. The qualitative enrichment profile is consistent with the ConsangBP positive control. Conversely, ARLQ genes showed no significant deviation from the fraction of the genomic background under strong recessive selection (OR=0.76, p=0.06), suggesting the prevalence of essentially random annotation errors. The fact that this gene set is fourfold the size of ARHQ suggests this is not due to a lack of statistical power. For comparison, we repeated this analysis for CGD AD and similarly found that the ADLQ set is indistinguishable from the genomic background with respect to depletion for strong recessive selection (OR=0.74, p=0.14), demonstrating this annotation quality issue is independent of MOI. In all cases the test for enrichment of strong additive selection is in the expected direction and significant, though only marginally in the case of ADLQ (ADHQ: OR=2.52, p=1.1 × 10^−8^; ADLQ: OR=1.28, p=0.022).

Last, we assessed the dependence of enrichment for strong recessive selection on the severity of autosomal recessive disease. We analyzed a list of effectively lethal, recessive Mendelian disease genes from a study of the population frequencies of such diseases [33]. This Lethal ARHQ list is a subset of the CGD ARHQ gene set that includes only the most extreme phenotypes leading to death before reproductive age, infertility in both sexes, or severe mental or physical developmental delay precluding reproduction. As seen in **Figure 4B**, this list of 23 scored genes, while only marginally significant due to the set size, showed a larger odds ratio than any other gene set tested (OR=2.92, p=0.042). This suggests that variants associated with effectively lethal recessive phenotypes may be identifiable through the action of recessive selection on their frequency distribution at the population level. The fact that an enrichment is observed in such a small gene set provides something of an empirical power bound on the minimal gene set size for aggregate detection of recessive selection.

To illustrate the more extensive space of analyses performed, we created `srMLGenes`, a web-based interactive tool that can be accessed at https://jordad05.u.hpc.mssm.edu/srmlgenes/ or downloaded at https://github.com/rondolab/srmlgenes/. This allows for full exploration of each of the aforementioned gene sets, as well as the ability to upload a novel gene set for the detection of enrichment for strong recessive selection (see **Supplementary Note 1**).

## Discussion

Here we present work aimed both at placing bounds on the power to infer recessive natural selection from large-scale human population data and developing a tool to aid in the identification and quantification of recessivity in the human genome. Few studies to date have inferred the presence of recessive selection from human population data [11, 18], yet many studies have focused on identifying genes with a recessive mode of inheritance in human disease [2–6] or on differentiating properties of AR diseases from those with AD [34, 35]. The validation of likelihood-based tests as a tool to identify the presence of recessive genes allows us to apply population genetic inference to the prioritization of AR disease genes, complementing a long tradition of computational predictions of (heterozygote) selection applied to Mendelian disease gene discovery pipelines [6, 21, 27–29, 31, 36–39]. The fact that we recapitulate an expected enrichment for strong recessive selection in genes annotated as involved in recessive disease by applying a quality filter demonstrates the potential power of this approach. The failure to detect an enrichment before applying this filter strongly suggests that a large fraction of genes annotated as AR and used in clinical genomics show little evidence of strong recessive selection in aggregate and are consistent with noise.

We established an empirical lower bound on the size of a gene set needed to detect enrichment for strong recessivity in current human population samples. In the Lethal ARHQ gene set, we found a significant enrichment for strong recessive selection in a set of only 23 genes [33]. Due to the nature of the phenotypes caused by these genes—infertility, death before reproductive age, etc.—LOF variants in these genes have no known phenotypes in heterozygote (carrier) form and are thus expected to have both the strongest selection and be exactly in the *h* = 0 corner. It is therefore unrealistic to expect any smaller gene sets categorized by srML to produce significant enrichments, regardless of their content.

Our work suggests theoretical and empirical power limitations on both the length of a recessive genes needed to inform any nontrivial inference of recessive selection and the number of genes needed to detect the presence of recessive selection in aggregate via enrichment. Observation of significant enrichments establishes that the human genome contains a sufficient number of genes evolving under strong recessive purifying selection to allow for detection. We found broad consistency in expected directions when comparing candidate recessive genes in both panmictic and consanguineous population data, between predictions of both high and low levels of haploinsufficient activity and recessivity, and finally between disease mode of inheritance and the mode of selection. Lastly, we produced a publicly accessible user interface where unique gene sets can be uploaded to assess their enrichment for genes evolving under strong recessive selection.

## Methods

### Filtering of Variants

All population data analysis was performed on non-Finnish European (NFE) samples in the Exome Aggregation Consortium (ExAC) dataset [21]. Before constructing SFSs, we first restricted to single nucleotide variants with only one alternative allele. Next, we removed variants where fewer than 80% of ExAC NFE samples had a confident genotype call to remove sites with poor coverage or ambiguous sequences. We removed sites where either allele formed a CpG hypermutable context to ensure a relatively homogenous mutation rate and prevent complications in the inference method due to recurrent mutations. To determine which allele was ancestral, we retrieved the orthologous site in the panTro2 chimpanzee sequence from the Ensembl Compara database, https://useast.ensembl.org/info/genome/compara/index.html [40]. For variants where one of the human alleles matched the orthologous panTro2 sequence, we assumed that the chimpanzee reference allele was the ancestral allele. Variants where neither allele matched the panTro2 sequence or where no orthologous site in panTro2 could be found were discarded.

### Calculation of Mutational Target Size and Gene Length

To calculate mutational target size for each gene, we first retrieved ExAC coverage regions from the gnomAD browser, https://gnomad.broadinstitute.org/downloads#exac-resources [28]. We then enumerated all possible mutations within the coverage regions and determined which of these passed our filtration steps and met our definition for either LOF and damaging (missense mutations annotated “probably damaging” by PolyPhen2 [25]) or coding synonymous sites. Finally, we applied the method of Samocha et al. [41] to calculate local mutation rates for each site meeting these criteria, and summed these rates into combined mutational target sizes for each gene and functional category. We converted these target sizes to approximate gene lengths by dividing by 10^−8^, an approximation of the average per-base mutation rate in the human genome.

### Calculation of Population Genetics Scores

Scores displayed in Fig. 1 were calculated as follows.

- For *π*_*ns*_*/π*_*s*_, we first calculated *π*_*ns*_ and *π*_*s*_ as

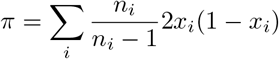

where *n*_*i*_ is the total number of observations made of allele *i* in the NFE subset of ExAC [21] (i.e. the allele number), and *x*_*i*_ is the frequency of that allele, summing over all synonymous (*π*_*s*_) or nonsynonymous (*π*_*ns*_) alleles in each gene, after filtering variants as described elsewhere. We removed all genes where either score was zero (that is, genes with no segregating sites in either functional class). We then ranked genes by |log *π*_*ns*_*/π*_*s*_| to measure deviation from a ratio of 1, and the highest 10% were counted as having evidence in favor of selection.

- For segregating sites, we ranked genes by number of sites with segregating alleles in the NFE subset of ExAC [21] divided by the calculated mutational target size after filtering variants as described elsewhere. The lowest 10% were counted as having evidence in favor of selection.
- For pLI, we retrieved pLI scores computed from gnomAD v2.1 for each gene covered in ExAC from the gnomAD browser, https://gnomad.broadinstitute.org/downloads [21, 28]. Genes with pLI scores greater than 0.9 were counted as having evidence in favor of selection.
- For O/E, we retrieved O/E loss of function scores from the same gnomAD v2.1 constraint data used for pLI [28]. We then ranked genes by O/E score. The lowest 10% were counted as having evidence in favor of selection.
- For RVIS, we retrieved the latest version of RVIS scores calculated from ExAC from http://genic-intolerance.org/data/RVIS_Unpublished_ExACv2_March2017.txt [27]. We then ranked genes by the standard
- RVIS score, labelled `(RVIS[pop_maf_0.05%(any)])`. The lowest 10% were counted as having evidence in favor of selection.
- For *s*_*het*_, we retrieved *s*_*het*_ estimates from Supplementary Table 1 of Cassa et al. 2017 [29]. We then ranked genes by estimated *s*_*het*_. The highest 10% were counted as having evidence in favor of selection.
- For dN/dS, we retrieved chimpanzee dN and dS values for each gene from version 88 of the Ensembl Biomart, https://www.ensembl.org/biomart/martview/ [19, 40]. We removed all genes where either score was zero (that is, genes with no substitutions in either functional class). We then ranked genes by |log dN*/*dS| to measure deviation from log [dN*/*dS = 1] = 0. The highest 10% were counted as having evidence in favor of selection.
- For phastCons, we retrieved phastCons scores for each site from the hg19 version of the UCSC genome browser, http://hgdownload.soe.ucsc.edu/goldenPath/hg19/phastCons100way/ [22, 42]. We calculated a per-gene phastCons score by averaging over the coding sequence of each gene. Genes with average phastCons scores greater than 0.9 were counted as having evidence in favor of selection.
- For the McDonald-Kreitman test [26], we first calculated *π*_*ns*_*/π*_*s*_ and dN/dS as described above. We then ranked genes by 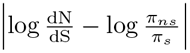 to measure deviation from a ratio of 1, and the highest 10% were counted as having evidence in favor of selection

In all cases, p-values were calculated using a chi-squared contingency test and confidence intervals were calculated assuming Poisson error on each cell of the contingency table.

### Forward Time Simulations

Genes with known selection and dominance were simulated through a realistic demography for the non-Finnish European population using a custom forward-time Wright-Fisher simulator implemented in Python, both of which were previously reported in Cassa, Weghorn, Balick, Jordan, et al. [29] and Weghorn, Balick, et al. [31]. The demographic history used was previously fit using the ExAC NFE population sample and corresponds to that of Tennesen, et al. with an increased rate of exponential growth during the most recent epoch [43]. We simulated a grid of 17 pairs of selection and dominance coefficients, comprising all pairwise combinations of *h* ∈ {0.0, 0.1, 0.3, 0.5} and *s* ∈ {−0.1, −0.01, −0.001, −0.0001} and one point at *s* = 0, which is identical for all values of *h*. To make reference simulations, we simulated 10^9^ unlinked sites with mutation rate *µ* = 10^−8^ per site per generation for each of the 17 selection and dominance classes. Sites were then binomially downsampled to a diploid sample size of 34,429, representing the sample size of NFE exomes used to calculate empirical site frequency spectra (SFS) from the ExAC database [21]. A reference SFS was calculated for each of the 17 selection and dominance classes using by counting the total number of alternate alleles in each frequency bin from the downsampled simulations. To make simulated genes, we simulated 10^6^ unlinked sites (again with *µ* = 10^−8^) for each of the 17 selection and dominance classes in the same way. We randomly sampled from these sites with replacement to create 31 simulated genes from each simulation on a logarithmically spaced grid ranging from 10^2^ to 10^5^ (i.e. 10^2^, 10^2.1^, 10^2.2^, …, 10^4.8^, 10^4.9^, 10^5^), corresponding to the approximate range of LOF and damaging single gene target sizes observed in the exome. We repeated this 10,000 times for each selection and dominance class, to produce a total of 10,000 simulated genes for each of 31 lengths and 17 selection and dominance classes. In total, we simulated 1.717 × 10^12^ unlinked sites and used them to create a reference SFS and 5.27 × 10^6^ simulated genes.

### Likelihood Calculation and Likelihood Tests

To compute the Poisson Random Field [30] likelihood of an observed SFS given a combination of dominance coefficient *h* and selection coefficient *s*, we compared the observed SFS with the SFS from a reference simulation of 10^9^ sites with the same *h* and *s* values. We first scaled the reference simulation SFS by the mutational target size *U*_*obs*_ for a gene with an observed SFS, so that it represents the expected SFS for a region with that target size:

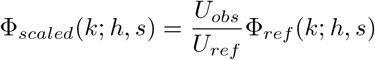

where *U*_*obs*_ and *U*_*ref*_ are the mutational target sizes corresponding to the observed and reference SFSs,and Φ_ref_(*k*; *h, s*) is the number of alleles seen in exactly *k* individuals in the reference simulation with dominance and selection coefficients *h* and *s*. For simulated genes and the reference simulations, these *U* values are exactly known; for empirical SFSs derived from ExAC, they were estimated as described above. Here we have assumed that the SFS is linearly scalable with mutational target size (i.e. the shape of the SFS is the same for all mutation rates). This is an approximation that comes from the infinite sites model and is predicated on linkage equilibrium and the absence of recurrent mutations. This approximation is invalid in the presence of substantial linkage disequilibrium or when the per population per site mutation rate is high enough to be observed in the sample. Notably, this restricts the method to panmictic population samples without recurrent mutations for the mutation spectra analyzed (i.e. non-CpG mutations). European samples larger than ExAC (*n* ≳ 10^4^) can be recurrent in most mutational classes, and the increased mutation rate of CpGs result in observed recurrent mutations even in ExAC [21].

Next we calculated the log likelihood under a Poisson Random Field (PRF) model [30]:

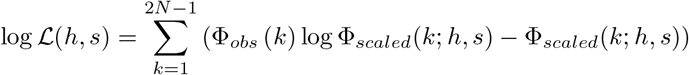

Where Φ_*obs*_(*k*) is the number of alleles seen in exactly *k* haploid chromosomes in the observed SFS, and *N* is the number of diploid individuals sampled.

For each observed SFS (i.e. for each gene and functional class), we performed this calculation once per reference simulation for a total of 17 combinations of *h* and *s*. This produces a log likelihood function log ℒ (*h, s*) defined at 17 discrete points. To perform the Likelihood Ratio Test (LRT) for non-additivity described in Williamson et al. [17], we applied a standard chi-squared likelihood-ratio test on this log likelihood function with the set of *h* = 0.5 points (including *s* = 0) as the null hypothesis. To perform the strong recessive maximum likelihood (srML) test, we partitioned the (*h, s*) plane into a strong recessive region consisting of the single simulated diploid selection class (*s* = −0.1, *h* = 0.0) and a non-strong-recessive region consisting of the other 16 diploid selection classes. We classify genes as “strong recessive” if the point that maximizes the PRF likelihood is (*s* = −0.1, *h* = 0.0), such that the log likelihood ratio is greater than 0, and as “not strong recessive” otherwise. An analogous test can also be used for other values of *h* and *s* where a different point in (*h, s*) space is compared to the remaining coordimates. In the text and figures, we use “strong recessive” to refer to the test described here for (*h* = 0.0, *s* = −0.1), and use “strong additive” to refer to the analogous test for (*h* = 0.5, *s* = −0.1) and “neutral” to refer to the analogous test for *s* = 0.

### Simulating Genomic Backgrounds

First, we pooled sites across all LOF and damaging sites in the ExAC NFE data to assemble an SFS, removing CpG and low coverage sites as described above [21]. Following the procedure detailed in the Supplement Information of Do et al. [44], we modeled an additive DFE by using a linear combination of simulated SFSs with three discrete selection strengths at (*h* = 0.5, *s* = {−10^−1^, −10^−2^, −10^−5^}) representing strong selection, weak selection, and near neutrality, respectively. We then estimated the maximum likelihood linear combination from a grid of three possible weights in steps of 0.05, where all weights sum to one for a total of two degrees of freedom. The PRF likelihood was maximized with the following weights: 0.6 for strong selection (*s* = −10^−1^), 0.3 for weak selection (*s* = −10^−2^), and 0.1 for nearly neutral sites (*s* = −10^−5^). As a control, this procedure was repeated for ExAC NFE synonymous sites with the same filtering steps, which resulted in 100% nearly neutral sites.

We then used this additive inference to simulate plausible genome wide distributions by assuming that the strong selection class inferred under additivity likely corresponds to strong additive selection, modeled as (*s* = −0.1, *h* = −0.5) for the present purposes, while the nearly neutral class can be treated as effectively neutral (*s* = 0, *h* = undefined) regardless of the additivity constraint. The proportion of the DFE associated with the weak selection class (30% attributed to *s* = −10^−2^) was decomposed into two discrete cases for simplicity: weak additive selection (*h* = 0.5, *s* = −10^−2^) and strong recessive selection (*h* = 0, *s* = −10^−1^). The fraction of this middle class was varied from 0 to 100% strong recessive (corresponding to an empirically derived maximum of 30% strong recessive selection and 0% weak additive in the genome) creating a range of possible genome wide joint DFEs. We matched the distribution of gene target sizes to that of LOF and damaging variants in ExAC NFE after the filtering steps described above (see **Figure SI1**). We simulated the 17 point grid of selection and dominance coefficients detailed above over a range of mutational targets (i.e. gene lengths) corresponding to the LOF and damaging range of genes in the exome: *L*_*LOF* +*damaging*_ ∈ [10^0.3^, 10^4.7^], where *L* = *U/*10^−8^, or between roughly 2 and 50000 bases. Using these simulations, we created an ensemble of genome-wide backgrounds, each with 14326 total genes, with a strong additive fraction = 0.6 and neutral fraction = 0.1 (together comprising 70% of genes), and with a strong recessive fraction = *f*_*R*_, and weak additive fraction = (0.3 - *f*_*R*_) (together comprising the remaining 30% of genes). Here *f*_*R*_ corresponds to the fraction of the genome that is under strong recessive selection up to a maximum of 0.3, the empirically estimated sum of the weak additive and strong recessive classes. We then simulated gene sets of various sizes (n= {30,100,300,…}) each with a known odds ratio from a depletion of OR=0.25 to an enrichment of OR=4.0 of recessive strong selection genes, with the remaining genes being pulled randomly from a genomic background with a given fraction *f*_*R*_.

### Gene Set Enrichment Analysis

For both simulated and empirical gene sets, we tested for enrichment of strong recessive selection against the genome-wide background using a chi-square contingency test. We constructed a contingency table where one variable represented membership in the gene set, and the other represented whether or not the gene was identified as strong recessive by the srML test. An enrichment (odds ratio *>*1.0) represents a scenario where genes in the gene set are more likely to be identified as strong recessive than genes not in the gene set, while a depletion (odds ratio *<*1.0) represents a scenario where genes in the gene set are less likely to be identified as strong recessive than genes not in the gene set. The test was performed as a two-talied test.

For the empirical gene sets, we also used logistic regression to attempt to control for gene length / mutational target size, which may be a confounder. We used gene set membership as a binary dependent variable, and the binary result of the srML test and the LOF and damaging mutational target size of the gene as independent variables. The results of this analysis were qualitatively similar to those of the chi-square test, and the effect size of the length variable found by regression was uniformly zero.

### Empirical Gene Sets

The putatively recessive-enriched ConsangBP gene set was derived from sequencing of a British Pakistani population with a high degree of consanguinity. Genes were collected from Supplementary Data S1 of Narasimhan et al. 2016 [11], and consist of all genes harboring rare homozygous loss-of-function variants in this population. While genes containing rare homozygous loss-of-function variants in a highly consanguineous population are not *a priori* subject to recessive selection, the high consanguinity of the cohort allowed the authors to measure recessive selection by comparing loss-of-function variants to frequency-matched synonymous variants, and this specific list of genes was found to be subject to significant recessive purifying selection.

The putatively recessive-depleted HI80 gene set was derived from the haploinsufficiency scoring method of Huang et al. 2010 [24]. This method gives genes a score representing the posterior probability of haploinsufficiency based on biological similarity to known haploinsufficient genes. We defined our HI80 gene set using genes scored with a posterior probability for haploinsufficiency of at least 0.8 by this method. The HI20 gene set was derived from the same method using genes scored with a posterior probability of haploinsufficiency no greater than 0.2.

Clinical gene sets were collected from the Clinical Genomic Database (CGD) [7]. Autosomal recessive (AR) disease genes were defined as genes containing known AR disease variants and no known AD or other classes of variants. Autosomal dominant (AD) disease genes were defined as genes only containing known AD disease variants and no AR or other classes of variants. We removed all genes located on sex chromosomes, as well as any gene with a mixed MOI annotated with both AR and AD variants (AR/AD genes) and/or imprinting.

The Lethal AR gene set was collected from Table S2 of Amorim et al. 2017 [33], listing a manually curated set of variants confidently known to cause a severe form of lethal, Mendelian, recessive disease.

To assemble high-quality (HQ) and low-quality (LQ) sets from the CGD AR, CGD AD, and Lethal AR sets, we used ClinVar variant annotations [32]. HQ genes are those for which ClinVar contains at least two variants whose review status is 2 stars (“multiple submitters, no conflicts”) or better. LQ genes are the complement of this set, those for which ClinVar contains 1 or fewer variants whose review status is 2 stars or better.

## Acknowledgements

The authors thank Ivan Adzhubei for generous assistance with empirical target size estimates. RD is supported by R35GM124836 from the National Institute of General Medical Sciences of the National Institutes of Health, R01HL139865 and R01HL155915 from the National Heart, Lung, and Blood Institute of the National Institutes of Health. SS is supported by R35GM127131 from the National Institute of General Medical Sciences, R01MH101244 from the National Institute of Mental Health, and R01HG010372 from the National Human Genome Research Institute of the National Institutes of Health. DMJ is supported by T32HL00782 from the National Heart, Lung, and Blood Institute of the National Institutes of Health. All content is solely the responsibility of the authors and does not necessarily represent the official views of the National Institutes of Health.

## Author contributions

DJB, DMJ, SRS, and RD conceptualized the study and designed the methodology. DJB and DMJ performed data curation, formal analysis, software development, visualization, and writing of the initial draft. SRS and RD supervised all analyses and reviewed and edited the text.

