## Supplemental Information for "Overcoming constraints on the detection of recessive selection in human genes from population frequency data"

Daniel J. Balick, Daniel M. Jordan, Shamil Sunyaev, and Ron Do

#### Table of Contents


##### Supplemental Notes

**Note SN1**: Supplemental Note on using `srMLgenes`, an interactive tool to explore enrichment patterns of strong recessive selection.

##### Supplemental Tables

**Tables SI1** through **SI3** on gene set enrichment statistics supplementing **Figure 4**.

##### Supplemental Figures

**Figures SI1** through **SI7** supplementing **Figure 2** and **Figure 3**.

##### Supplemental Files

**File SF1**: Excel spreadsheet containing gene names, target sizes per functional class, PRF maximum likelihood $h$ and $s$ values, and results of the srML test for all genes with segregating sites and sufficient coverage in the non-Finnish European sample of the Exome Aggregation Consortium dataset.

### Supplementary Note on srMLgenes

#### An interactive tool to identify enrichment for strong recessive selection

To enable deeper exploration of the data presented here, we have developed `srMLGenes`, a web-based tool that allows interactive plotting of our data. This tool can be accessed at `https://jordad05.u.hpc.mssm.edu/srmlgenes/` or downloaded at `https://github.com/dmjordan/heatmaps_wsgi` and run on a local machine. This Supplementary Note outlines some of its features. There are two major components to the online tool: exploration of simulated genes and exploration of real human genes observed in ExAC data [1]. These are represented as two tabs at the top of the page.

##### Simulated Genes

The "Simulated Genes" tab shows a 2-dimensional histogram of the maximum likelihood selection and dominance classes for a subset of the 5,270,000 genes simulated for our analysis. This is represented as a heatmap over a grid spanning the 17 selection and dominance classes we used in our simulations (**Figure SN1.1**). The $x$-axis displays the selection coefficient, $s$, with strong selection ($s = -10^{-1}$) on the right and neutrality ($s = 0$) on the left. The $y$-axis displays the dominance coefficient, $h$, with recessivity ($h = 0$) on the bottom and additivity ($h = 0.5$) on the top. The leftmost column represents neutrality, for which $h$ is undefined, so all information about neutrality is contained in the top left corner and the grey cells should be ignored.

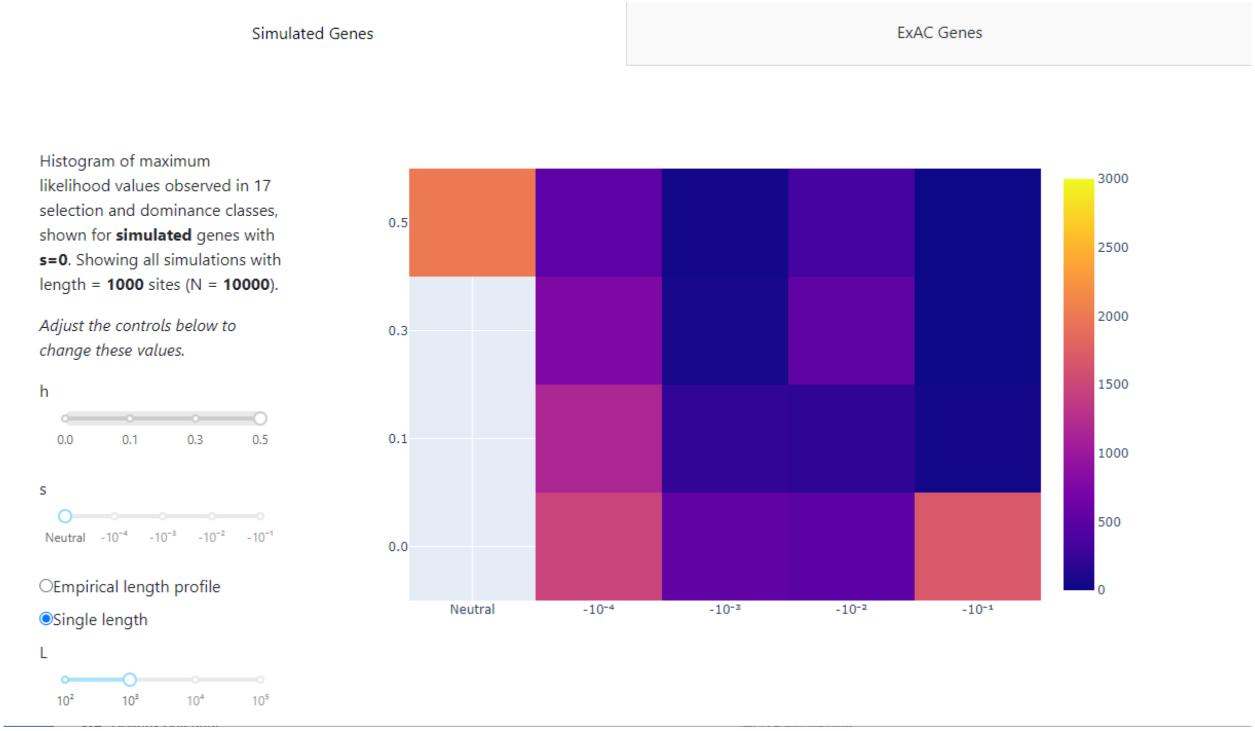


**Figure SN1.1:** The web interface when first opened, showing a histogram of simulated genes.

On the left side of the screen is a panel allowing the user to select the subset of simulations plotted (**Figure SN1.2**). The user can select a single combination of $h$ and $s$ values using the sliders labelled "h" and "s." When the "single length" button is selected, the user can select a simulated length using the slider labeled "L." The heatmap will then show the distribution of all 10,000 simulated genes with that combination of $h$, $s$, and length (**Figure SN1.2A**). When the "empirical length profile" button is selected, the user can select a gene list to use as a length profile (**Figure SN1.2B**). The heatmap then shows the distribution of a random sample of simulated genes that matches the number and lengths of the selected gene list. Note that the actual SFSs of these genes are not used in any way, only their length distribution. For details on the gene list selection interface, see **Gene List Selection** below.

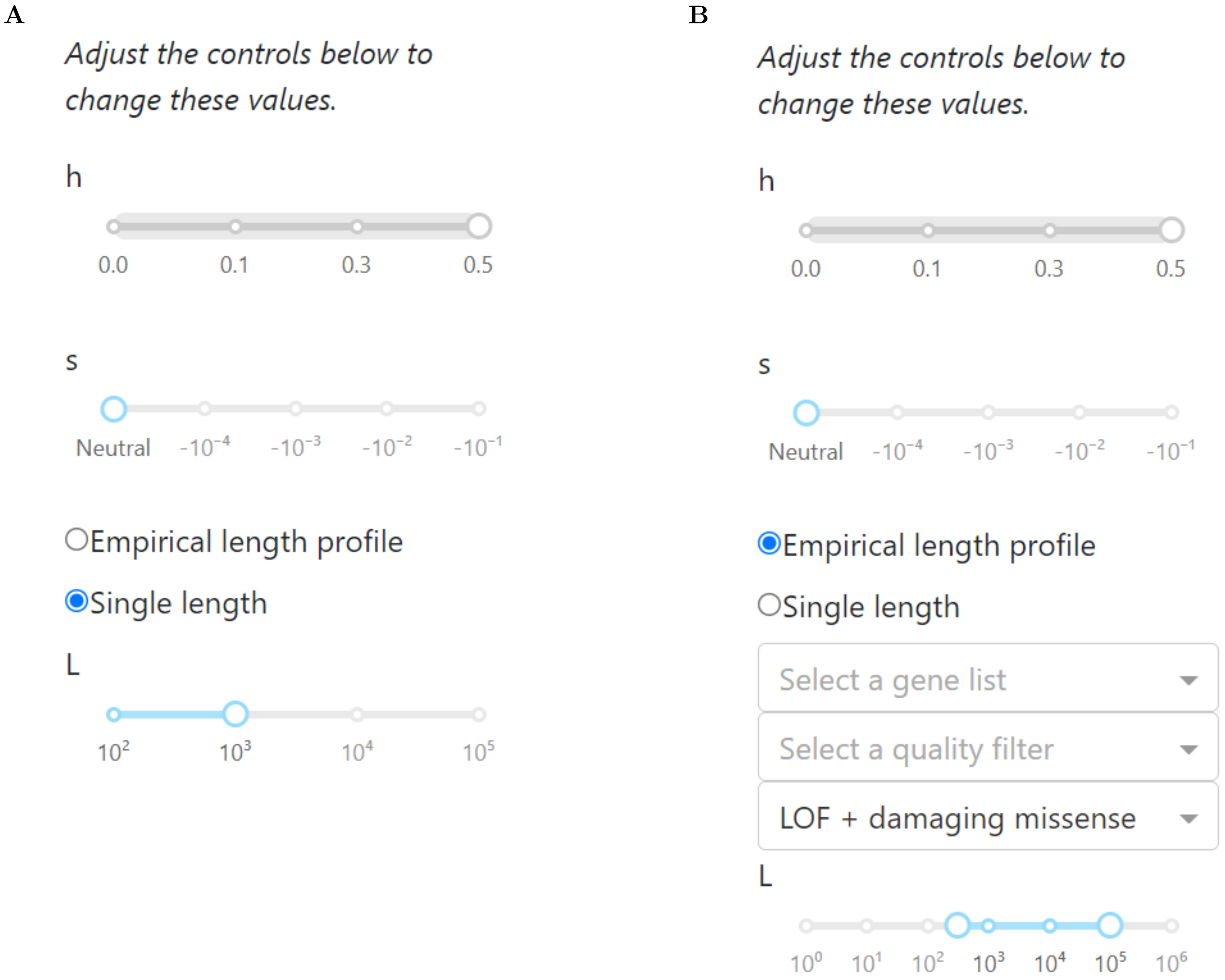


**Figure SN1.2:** Detail of the control panel allowing selection of a subset of simulations with a single $h$ and $s$. The slider for $h$ is grayed out when $s$ is set to neutral, since $h$ is undefined under neutrality. **A.** The control panel when "Single length" is selected, allowing selection of a single value for $L$. **B.** The control panel when "Empirical length profile" is selected, allowing selection of a gene set to base the simulation length profile on.

For either choice, a description of the currently displayed figure is shown to the left of the plot, above the controls, and hovering the mouse over each cell of the plot displays a tooltip with more detailed information about the individual selection and dominance classes (**Figure SN1.3**).

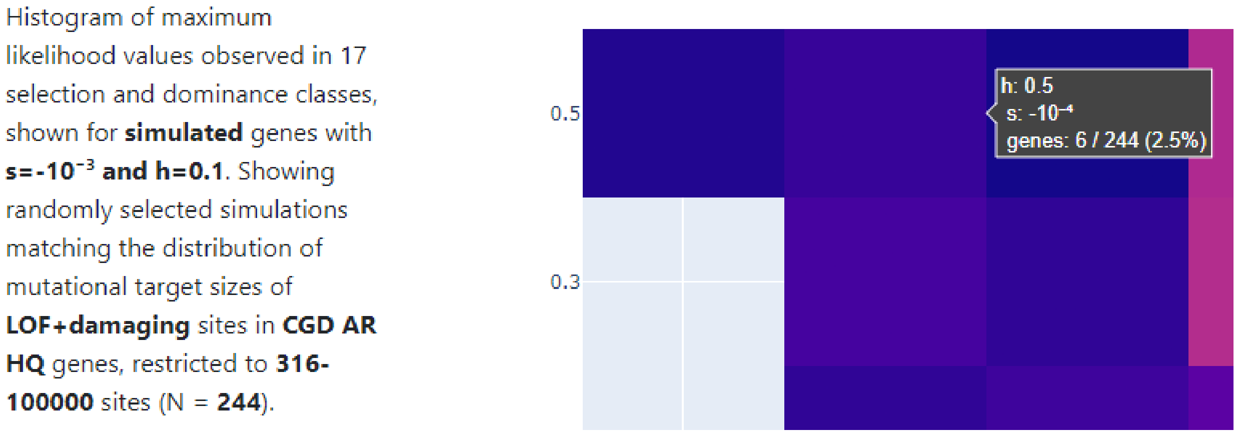


**Figure SN1.3:** Detail of the dynamically updating caption and mouseover tooltip for an example parameter choice.

##### ExAC Genes

Selecting the "ExAC Genes" tab reveals a slightly different set of controls (**Figure SN1.4**).

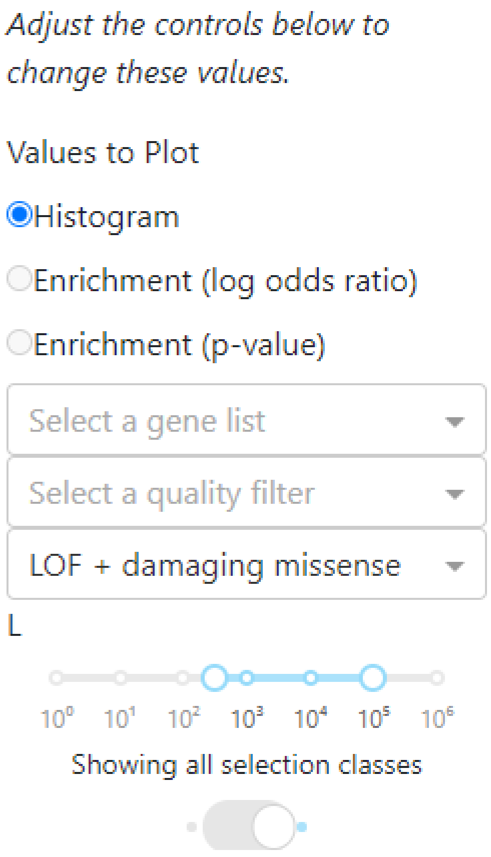


**Figure SN1.4:** Detail of the control panel for plotting ExAC genes. The radio buttons for enrichment plots is grayed out when no gene list is selected, since enrichment does not make sense without a specific gene list.

The ExAC tab can show one of three kinds of plots. When “Histogram” is selected, it shows a histogram of maximum likelihood values, similar to the “Simulated Genes” tab (**Figure SN1.5**)

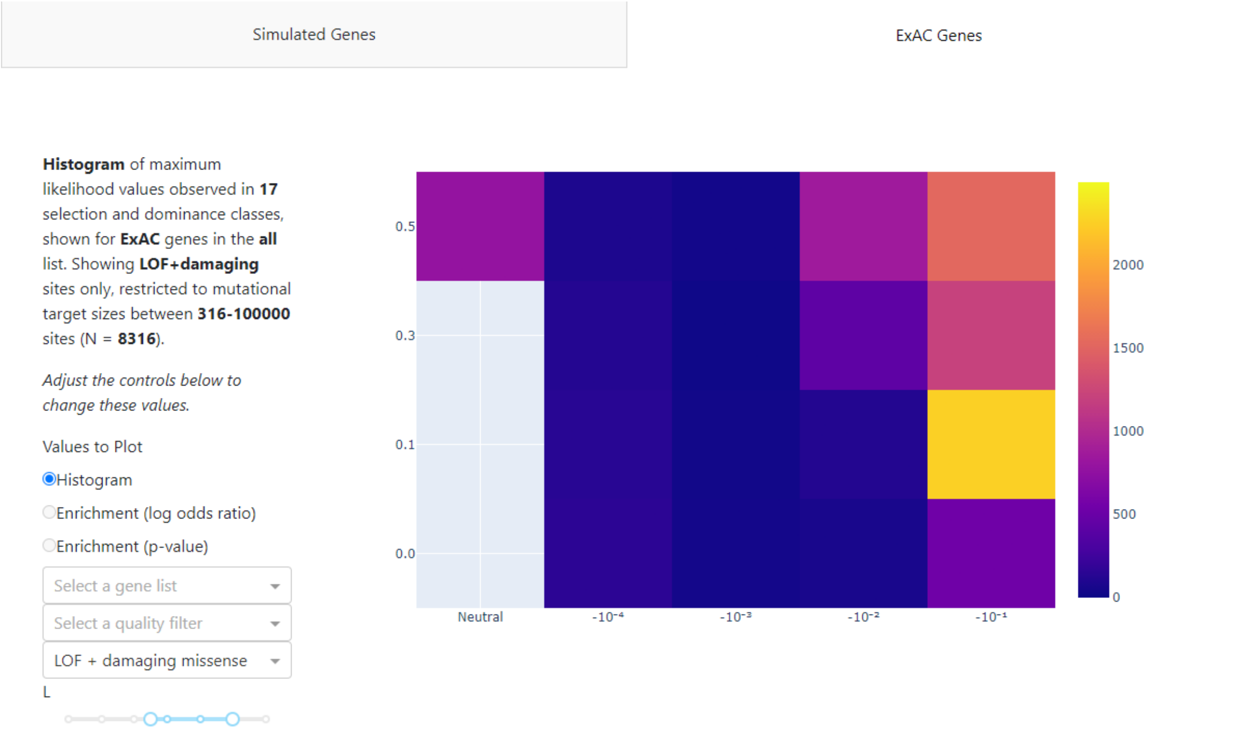


**Figure SN1.5:** The ExAC Genes tab when first opened, showing a histogram of all genes within the default length range.

When a gene list has been selected, “Enrichment (log odds ratio)” and “Enrichment (p-value)” also become available (**Figure SN1.6**). These options show the enrichment or depletion in each selection and dominance class for the selected gene list. In either case, enrichment is represented with blue and depletion with red.

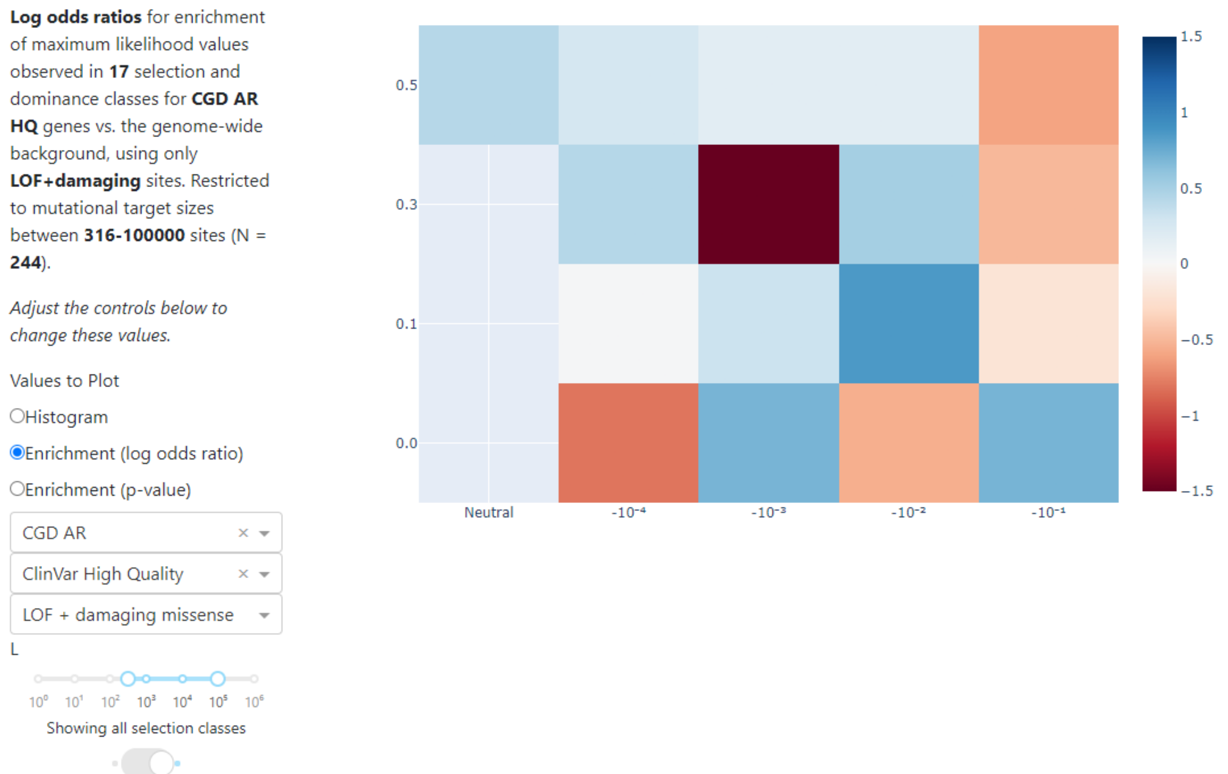


**Figure SN1.6:** Enrichment heat map view, showing a heatmap of enrichment in each selection and dominance class for CGD ARHQ genes.

There is also a switch at the bottom of the panel that toggles between "all selection classes" and "only strong selection" (it may require scrolling down to see this switch). When "all selection classes" is selected, the interface shows a heatmap, similar to the "Simulated Genes" tab. When "only strong selection" is selected, the interface shows a bar chart for the two selection classes we focused on in our analysis: strong additive ($s = -10^{-1}, h = 0.5$) and strong recessive ($s = -10^{-1}, h = 0.0$) (**Figure SN1.5**). This provides a simplified view to make exploration of results more intuitive (**Figure SN1.7**).

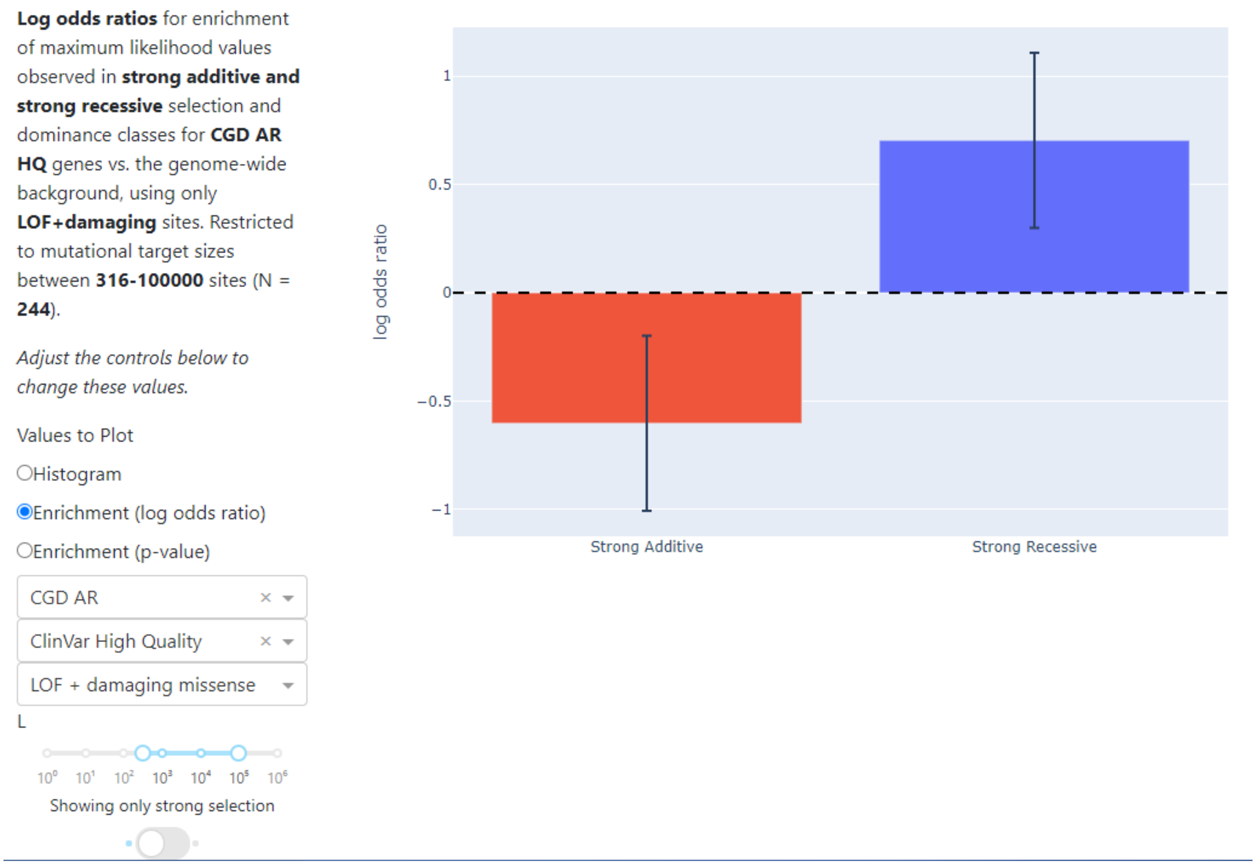


**Figure SN1.7:** Enrichment bar chart view, showing a bar chart of enrichment in the strong additive and strong recessive classes for CGD ARHQ genes.

As in the simulation tab, mousing over individual cells or bars shows detailed information about each cell, including the numerical enrichments and p-values (**Figure SN1.8**).

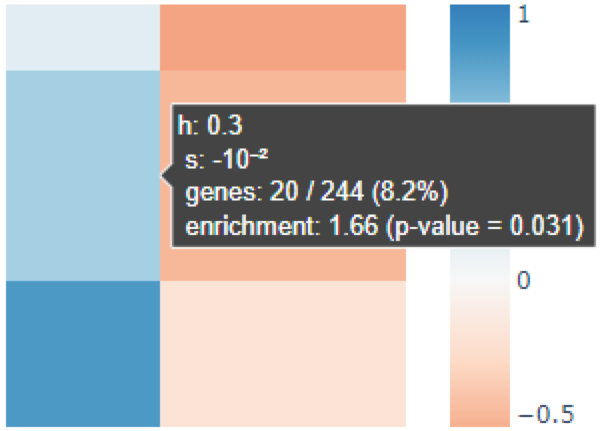


**Figure SN1.8:** Detail of a mouseover tooltip on the enrichment heat map view. Similar tooltips appear in the histogram view and the enrichment bar plot view.

##### Gene List Selection

In both tabs, a series of 4 controls allows users to specify a gene list (**Figure SN1.9**).

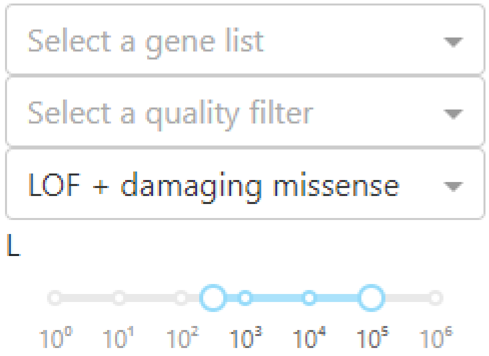


**Figure SN1.9:** Detail of the gene list specification interface as it appears when first opened.

The first control is the gene list drop down. This allows users to select one of the 6 gene lists analyzed in our manuscript (see **Figure 4**): ConsangBP, HI80, HI20, CGD AR, CGD AD, and Lethal AR (**Figure SN1.10A**). If it is left blank (the default), all genes are used. In addition to the gene sets used for our analysis, `srMLGenes` allows users to upload a custom gene list. If "Upload custom list" is selected, a text box and an upload button become available (**Figure SN1.10B**). Genes can either be entered into the text box or uploaded as a text file. In either case, uploaded genes should be in the form of HGNC gene symbols, separated by commas, spaces, or line breaks. Gene symbols that are not recognized will be ignored and removed from the analysis. After a gene list is either read from the text box by pressing the "Update" button or uploaded by pressing the "Select File" button, the total number of recognized genes from a custom gene list is reported, to help diagnose problems with unrecognized symbols (**Figure SN1.10C**). An exhaustive list of gene symbols recognized by `srMLGenes` can be found in the "gene" column of **Supplementary Data File 1**.

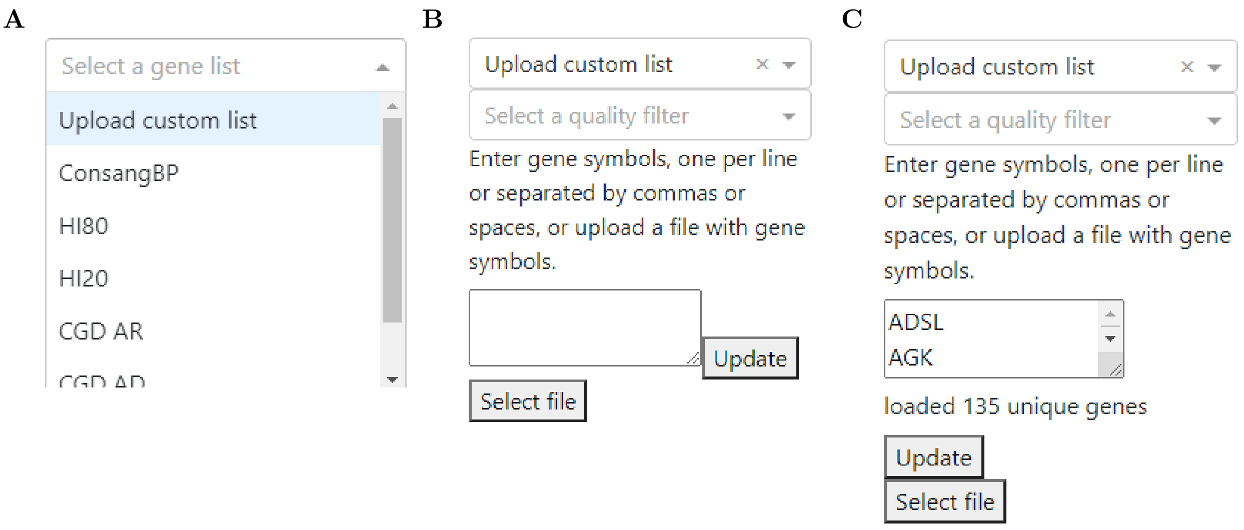


**Figure SN1.10:** Gene list dropdown interface. **A.** The dropdown list offers a choice of predefined gene lists as well as an "Upload custom list" option. **B.** A text box and upload button appear when "Upload custom list" is selected. **C.** The interface after a custom gene list has been loaded.

The second control is the quality filter drop down (**Figure SN1.11A**). If "ClinVar High Quality" is selected, the gene list is filtered to remove all genes containing fewer than 2 variants with ClinVar review status of 2 stars or/ higher, the same analysis used to produce the HQ gene lists shown in **Figure 4B** of our manuscript. If "ClinVar Low Quality" is selected, the gene list is filtered to remove all genes containing more than 1 variant with ClinVar review status of 2 stars or higher, the same analysis used to produce the LQ gene lists. If it is left blank (the default), no additional filtering is performed. Note that applying the ClinVar High Quality filter on gene lists that are not derived from monogenic disease data, especially for custom uploaded lists, may result in most of the genes being filtered out.

The third control is the function drop down (**Figure SN1.11B**). If "LOF+damaging missense" is selected (the default), the gene lengths and SFSs used are for loss of function variants and missense variants predicted to be damaging (classified as"probably damaging" by PolyPhen2 [2]), as discussed in our manuscript. If "synonymous" is selected, the gene lengths and SFSs used are for synonymous variants only. Synonymous enrichment patterns can act as a negative control in some analyses, but should be interpreted cautiously, as coding synonymous variants are not necessarily expected to be strictly neutral.

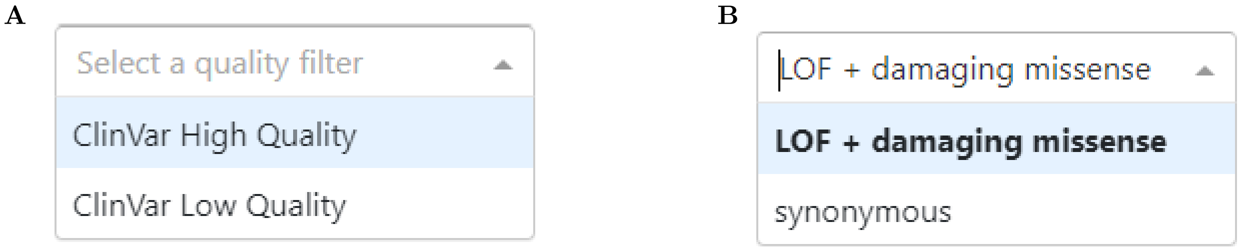


**Figure SN1.11:** Additional drop down menus for the gene list selection interface. **A.** The quality filter drop down, providing a choice of high quality, low quality, or no filter. **B.** The molecular function drop down, providing a choice of LOF + damaging missense or coding synonymous.

The last control is the length range. This allows specification of lower and upper bounds on gene length. The default is a range from $10^{2.5}$ (approximately 316 bases) to $10^5$, roughly the same as the range used in all analyses in our manuscript. (The lower bound differs slightly from the value of 300 used in the main text to preserve even spacing in $\log_{10}$ space.)

#### Supplementary Tables

**Table SI1:** Enrichment and depletion results for empirical gene sets using a chi squared contingency test.

| | | Strong recessive | | | Strong additive | | |
|---|---|---|---|---|---|---|---|
| **Gene Set** | **# scored** | **# ML** | **OR** | **p-value** | **# ML** | **OR** | **p-value** |
| ConsangBP | 401 | 40 (15%) | 2.60 | $1.9 \times 10^{-11}$ | 33 (8%) | 0.41 | $1.2 \times 10^{-7}$ |
| HI80 | 583 | 21 (4%) | 0.50 | 0.002 | 176 (30%) | 2.07 | $6.7 \times 10^{-15}$ |
| HI20 | 1170 | 122 (10%) | 1.77 | $7.0 \times 10^{-8}$ | 128 (11%) | 0.51 | $5.1 \times 10^{-12}$ |
| CGD AR | 1279 | 87 (7%) | 1.01 | 0.94 | 170 (13%) | 0.65 | $9.3 \times 10^{-7}$ |
| CGD ARHQ | 255 | 32 (13%) | 2.04 | $1.8 \times 10^{-4}$ | 28 (11%) | 0.55 | 0.0025 |
| CGD ARLQ | 1024 | 55 (5%) | 0.76 | 0.06 | 142 (14%) | 0.70 | $1.4 \times 10^{-4}$ |
| CGD ADHQ | 161 | 4 (2%) | 0.35 | 0.03 | 57 (35%) | 2.52 | $1.1 \times 10^{-8}$ |
| CGD ADLQ | 520 | 27 (5%) | 0.74 | 0.14 | 114 (22%) | 1.28 | 0.022 |
| Lethal AR | 23 | 4 (17%) | 2.92 | 0.042 | 4 (17%) | 0.95 | 0.92 |

**Table SI2:** Enrichment and depletion results for empirical gene sets using logistic regression, with length as a potentially confounding factor.

| | Strong recessive | | Strong additive | |
|---|---|---|---|---|
| **Gene Set** | **OR** | **p-value** | **OR** | **p-value** |
| ConsangBP | 2.65 | $4.04 \times 10^{-11}$ | 0.38 | $2.60 \times 10^{-07}$ |
| HI80 | 0.51 | $3.10 \times 10^{-03}$ | 2.05 | $4.07 \times 10^{-14}$ |
| HI20 | 1.69 | $1.3 \times 10^{-6}$ | 0.52 | $1.53 \times 10^{-11}$ |
| CGD AR | 1.04 | 0.73 | 0.64 | $6.71 \times 10^{-07}$ |
| CGD AD | 0.71 | 0.076 | 1.54 | $3.24 \times 10^{-06}$ |
| CGD ARHQ | 2.11 | 0.00012 | 0.53 | 0.0020 |
| CGD ARLQ | 0.77 | 0.078 | 0.69 | 0.00012 |
| CGD ADHQ | 0.41 | 0.088 | 2.44 | $1.23 \times 10^{-07}$ |
| CGD ADLQ | 0.76 | 0.18 | 1.27 | $2.90 \times 10^{-02}$ |
| Lethal AR | 2.97 | 0.049 | 0.94 | 0.92 |

**Table SI3:** Effect of length on empirical gene set membership, as detected by logistic regression.

| | Strong recessive | | Strong additive | |
|---|---|---|---|---|
| **Gene Set** | **OR** | **p-value** | **OR** | **p-value** |
| ConsangBP | 1.00 | 0.066 | 1.00 | 0.078 |
| HI80 | 1.00 | 0.034 | 1.00 | 0.053 |
| HI20 | 1.00 | 0.0021 | 1.00 | 0.00042 |
| CGD AR | 1.00 | 0.0089 | 1.00 | 0.0060 |
| CGD AD | 1.00 | $8.30 \times 10^{-13}$ | 1.00 | $2.67 \times 10^{-13}$ |
| CGD ARHQ | 1.00 | 0.021 | 1.00 | 0.011 |
| CGD ARLQ | 1.00 | 0.12 | 1.00 | 0.065 |
| CGD ADHQ | 1.00 | $2.24 \times 10^{-12}$ | 1.00 | $7.11 \times 10^{-13}$ |
| CGD ADLQ | 1.00 | 0.030 | 1.00 | 0.035 |
| Lethal AR | 1.00 | 0.66 | 1.00 | 0.79 |

#### Supplementary Figures

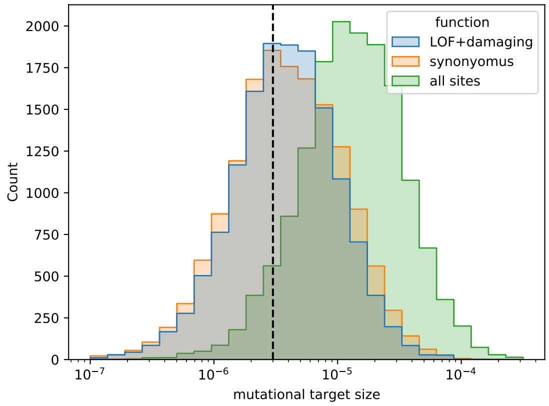


**Figure SI1:** Histogram showing the distribution of mutational target size per gene in the human exome. The dashed line indicates the length cutoff of $U \geq 3 \times 10^{-6}$ used for analyses.

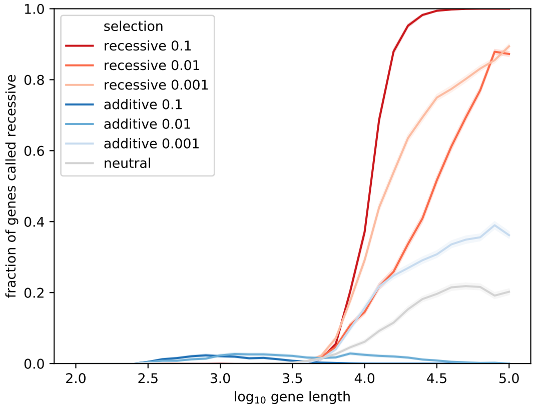


**Figure SI2:** Plot showing the fraction of simulated genes from each selection and dominance class rejecting the null hypothesis of additivity with LLRT, including weaker recessive selection.

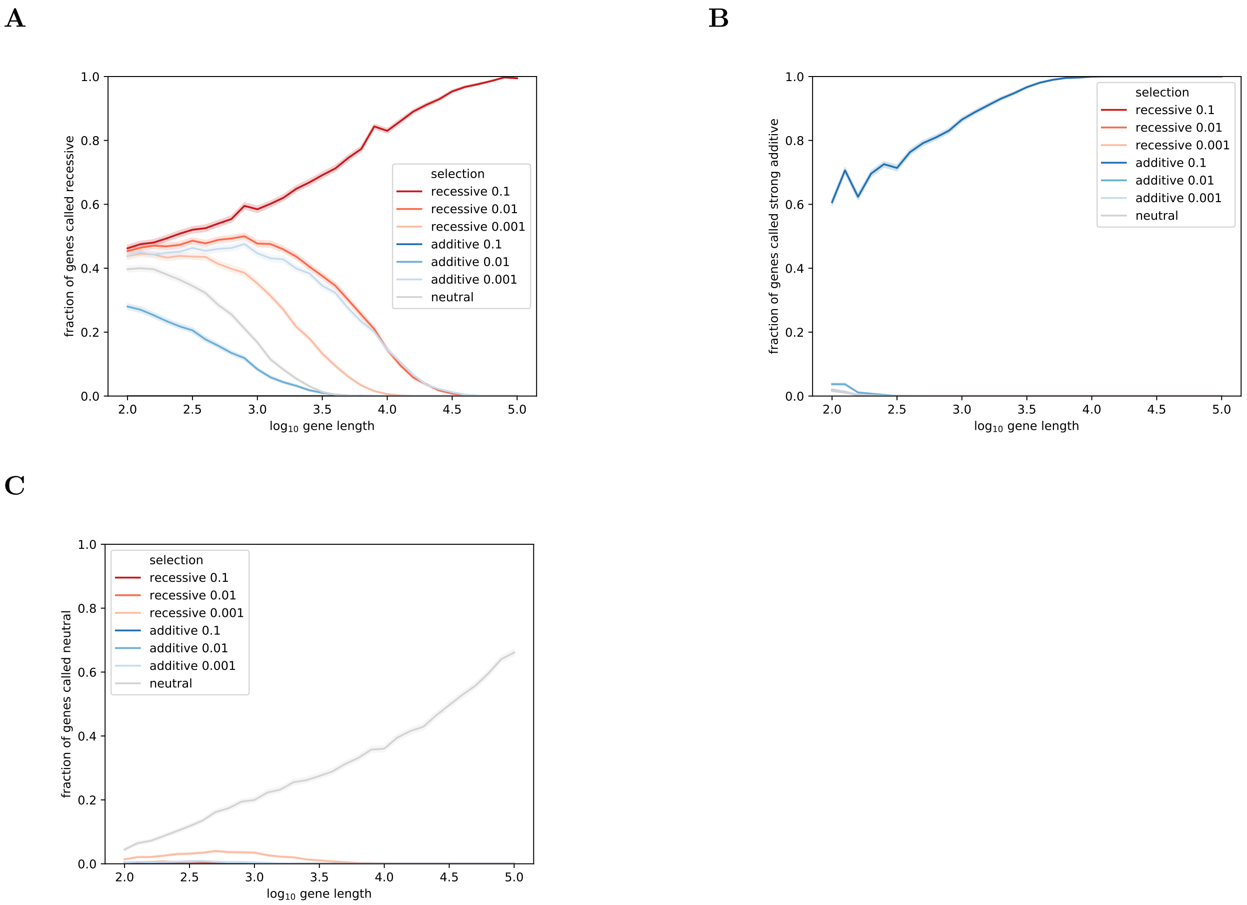


**Figure SI3:** The fraction of simulated genes with various selection and dominance coefficients identified by srML as strong recessive, strong additive, and neutral. Identification of the correct class (e.g. strong recessive selection correctly identified as strong recessive) corresponds to sensitivity of the test. Incorrect identification (e.g. weak additive selection misclassified as strong recessive selection) corresponds to the rate of false positives in the test due to genes with those specific selection and dominance coefficients. **A.** The fraction of simulated genes with various selection and dominance coefficients, including weaker recessive selection, identified as strong recessive by the srML test. Sensitivity is shown in red. **B.** The fraction of simulated genes with various selection and dominance coefficients identified as strong additive ($s = -0.1$, $h = 0.5$) by the strong additive equivalent of the srML test (i.e. a saML test). Sensitivity is shown in dark blue. Lines that are not visible are zero throughout. **C.** The fraction of simulated genes with various selection and dominance coefficients identified as neutral ($s = 0$) by the neutral equivalent of the srML test (i.e. an nML test). Sensitivity is shown in grey. Lines that are not visible are zero throughout.

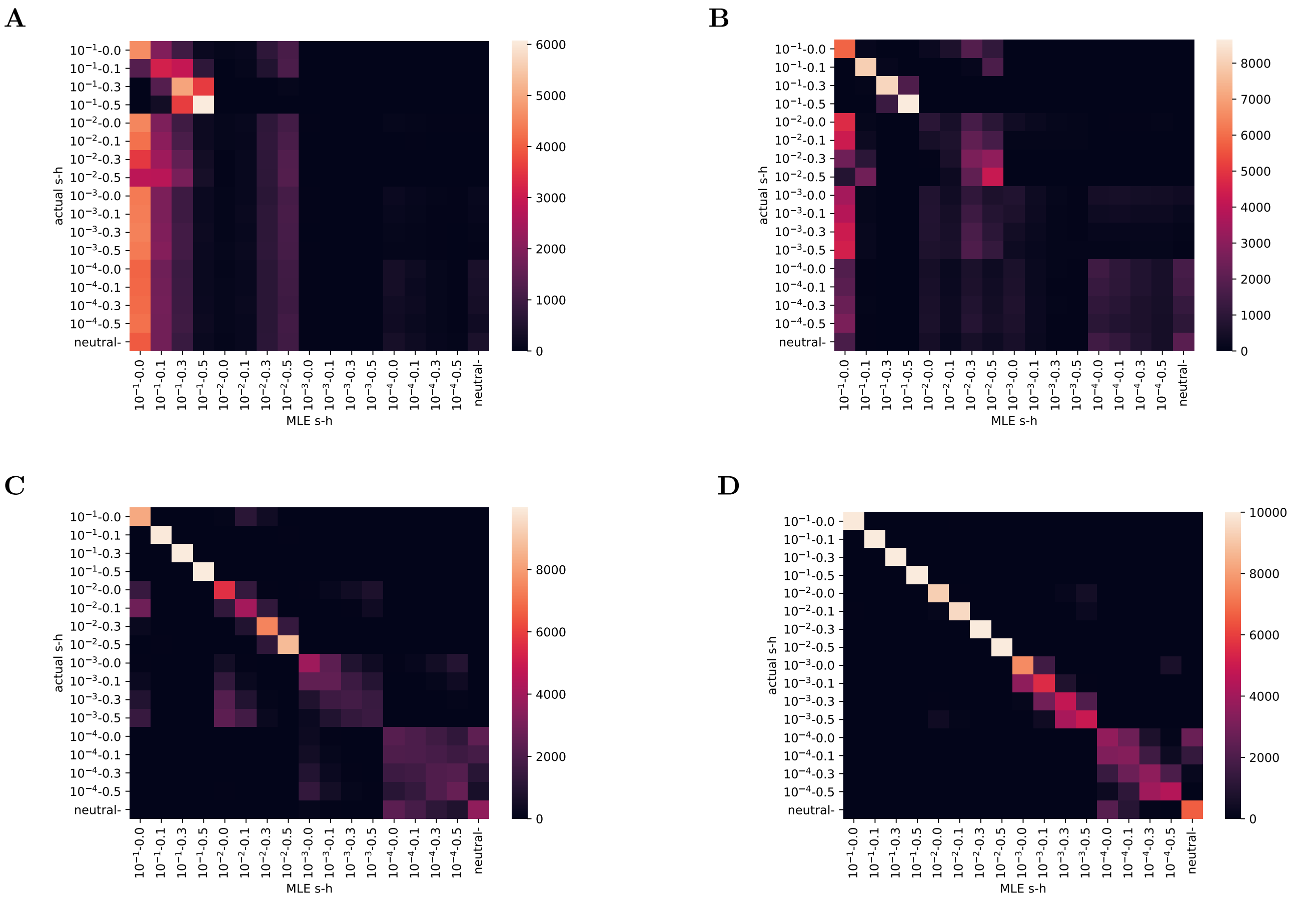


**Figure SI4:** Heatmap showing maximum likelihood estimates of selection and dominance for simulated genes of several specific lengths. $10^4$ simulated genes are shown for each $(h, s)$ coordinate such that each row sums to $10^4$ and that is the maximum value of any cell. Diagonal cells correspond to ML inference at the exact $(h, s)$ values simulated. **A.** Genes with length $10^2$ bp. **B.** Genes with length $10^3$ bp. **C.** Genes with length $10^4$ bp. **D.** Genes with length $10^5$ bp.

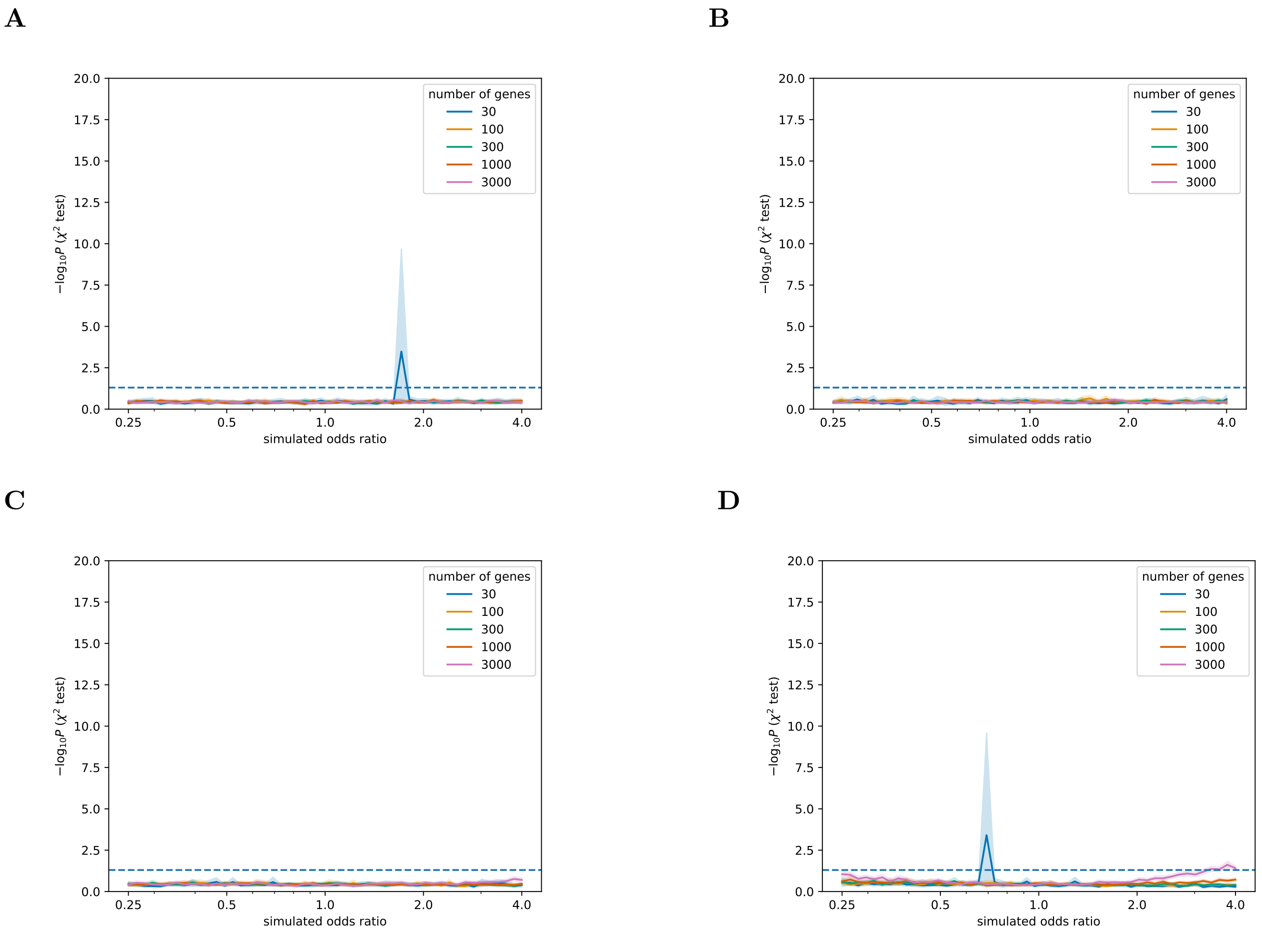


**Figure SI5:** Power to detect enrichment or depletion of genes rejecting the null hypothesis of additivity using the LLRT when a specific fraction of the genome is under strong recessive selection, $f_R$, an unknown quantity in humans. Power is shown for various gene set sizes in each plot. This test was found to be universally underpowered for all $f_R$ values consistent with empirical data. Spikes were not reproducible with additional replicates and should be interpreted as simulation stochasticity. **A.** $f_R$=0.1% of the genome under strong recessive selection. **B.** $f_R$=1% of the genome under strong recessive selection. **C.** $f_R$=10% of the genome under strong recessive selection. **D.** $f_R$=30% of the genome under strong recessive selection.

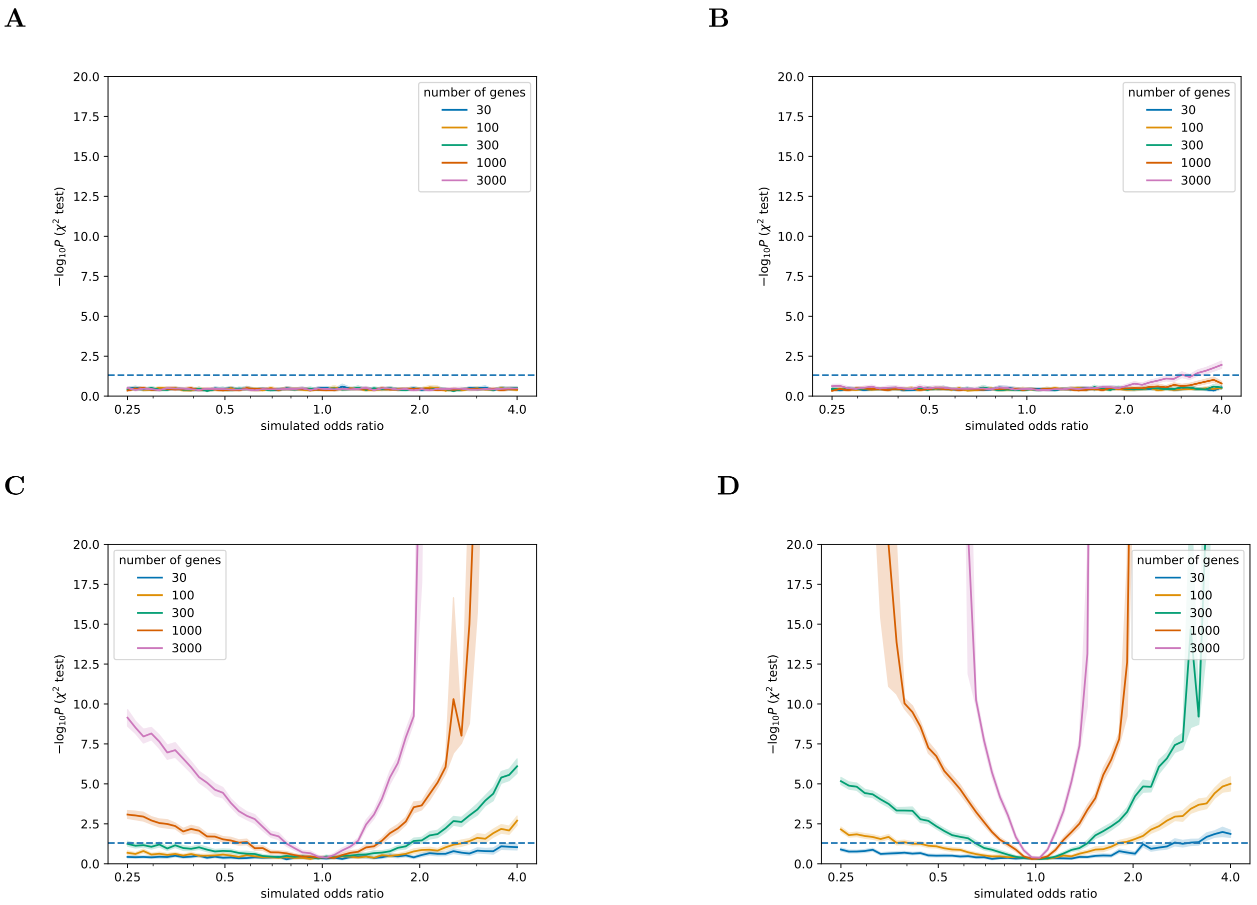


**Figure SI6:** Power of the srML test to detect enrichment or depletion for strong recessive selection when a specific fraction of the genome is under strong recessive selection, $f_R$, an unknown quantity in humans. Power is shown for various gene set sizes in each plot. When the $f_R$ is larger, this test has more power for any gene set size or odds ratio. The asymmetry of these plots shows that the srML test has more power to detect enrichment than depletion. **A.** $f_R$=0.1% of the genome under strong recessive selection. **B.** $f_R$=1% of the genome under strong recessive selection. **C.** $f_R$=10% of the genome under strong recessive selection. **D.** $f_R$=30% of the genome under strong recessive selection.

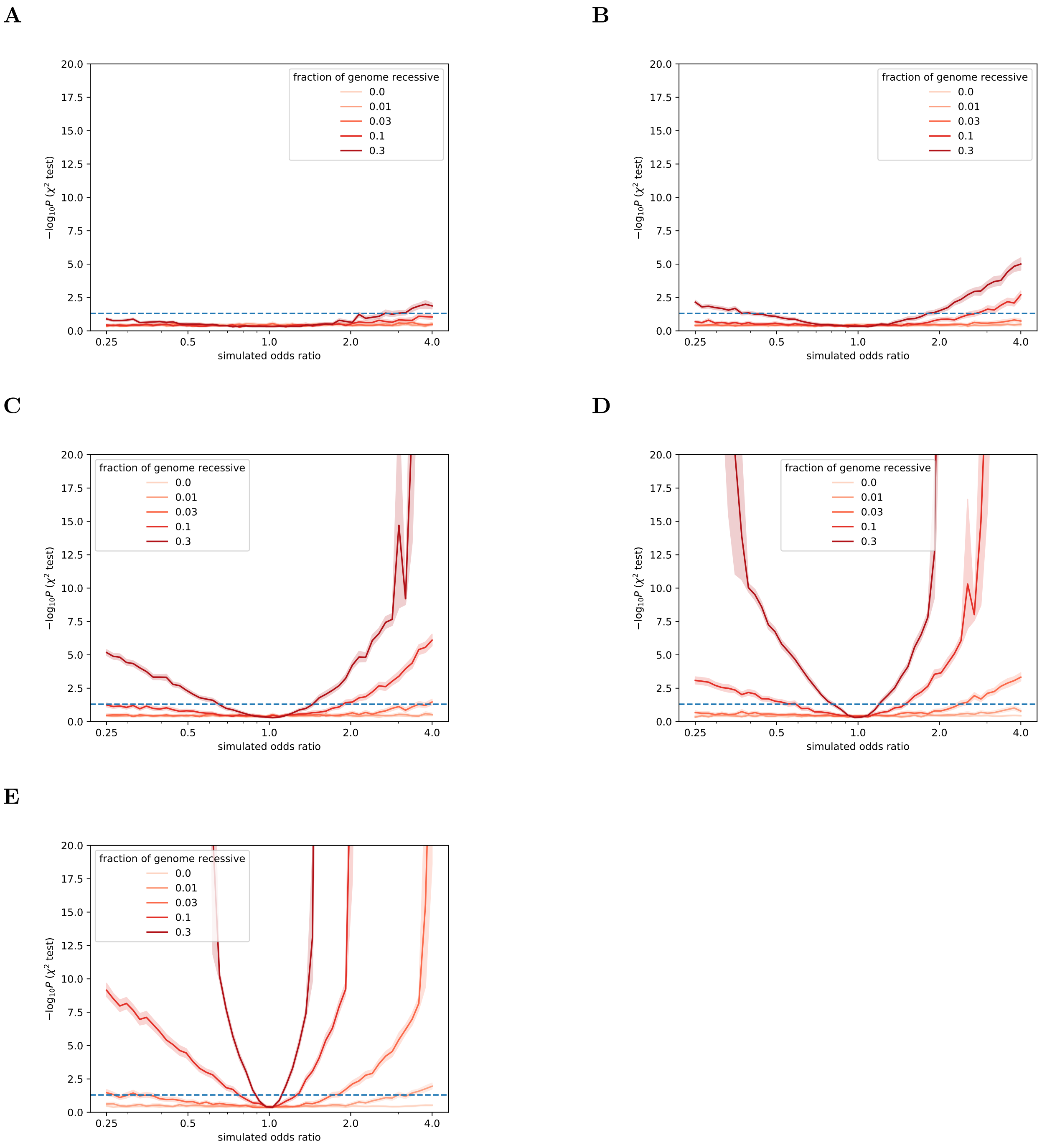


**Figure SI7:** Power of the srML test to detect enrichment or depletion for strong recessive selection for gene sets of a specific size. Power is shown for a range of fractions of the genome under strong recessive selection, $f_R$, in each plot. **A.** $N$=30 genes. **B.** $N$=100 genes. **C.** $N$=300 genes. **D.** $N$=1000 genes. **E.** $N$=3000 genes.
